# Surveying the Hormonome of Hazelnut Catkins During Winter Dormancy

**DOI:** 10.1101/2025.10.25.684543

**Authors:** John M. Steele, Sagar Datir, John K. Robinson, Sharon Regan

## Abstract

**Background:** Deciduous woody perennials, such as hazelnut, undergo winter dormancy to protect sensitive tissues, such as flowers, from harsh conditions. The reproductive success of the tree is dependent on the release of dormancy under favorable conditions. To bloom, the tree must first experience a certain amount of chilling, followed by a certain amount of warmth. With global warming, many trees risk not being able to accumulate enough chilling to release dormancy. Also, when trees accustomed to warm climates are brought into cold climates, they might bloom prematurely at the first sign of spring, and risk freezing damage. The latter is the case for hazelnut, recently adopted as a crop in Ontario, Canada. The present study investigates the hormonal regulation of dormancy in hazelnuts’ male flowers (catkins) by generating hormone profiles in early and late-blooming accessions throughout the dormant season. Abscisic acid (ABA), gibberellin (GA), auxin, cytokinin (CTK), their metabolites, as well as the ethylene precursor 1-aminocyclopropane-1-carboxylic acid (ACC), were measured.

**Results:** ABA decreased with dormancy progression, while GA increased. This correlation implies ABA is primarily responsible for dormancy maintenance in catkins and GA works antagonistically to ABA. Indeed, the ABA/GA ratio steadily decreased throughout dormancy. For the first time, CTKs have been reported to steadily increase during dormancy. Auxin and ethylene appear to primarily play a role in the onset of dormancy. Interestingly, early blooming accessions failed to accumulate the auxin conjugate, IAA-Asp and had higher ACC levels throughout most of dormancy.

**Conclusions:** Cumulatively, the present study has generated the most comprehensive hormone profile in dormant flowers of deciduous woody perennials and has identified potential strategies for the delay of bloom in hazelnut catkins through the manipulation of hormones.

## Background

Hazelnuts are nutrient rich, tasty nuts which have been a part of the human diet for millennia (Holstein et al., 2018). Most of the hazelnut cultivation, historically and currently, takes place in the Mediterranean basin (Boccacci & Botta, 2009). The hazelnut industry has recently expanded to southern Ontario, Canada (Dale et al., 2012). This expansion was primarily driven by the establishment of a hazelnut processing plant in Brantford, Ontario, by the Ferrero confectionery company (Dale et al., 2012). Hazelnuts are monoecious, dichogamous, self-incompatible, wind pollenated, winter-flowering trees (Botta et al., 2019). Successful nut production depends on overlap between pistil bloom of producers and pollen shed of compatible pollinizers. Male and female flowers begin developing late spring and early summer, respectively. Male flowers develop in cylindrical clusters called catkins. Catkins elongate as they develop, reaching a plateau in the fall in preparation for dormancy, at which time they are fully developed. Once dormancy is released, catkins open their flowers, elongate rapidly, and shed their pollen. In contrast to male flowers, female flower buds are not fully developed by the time of dormancy establishment, and are visually indistinguishable from vegetative buds until the release of dormancy (Olsen, 2024).

In conventional growing regions, hazelnuts are protandrous, with the catkins blooming before female flowers (Capik & Molnar, 2014). In regions with long, cold winters, however, hazelnuts tend to be protogynous, with female flowers blooming first (Capik & Molnar, 2014). Flowering typically takes place from December to March in the northern hemisphere, with peak bloom varying each year depending on local weather patterns (Capik & Molnar, 2014). Catkins typically exit this state of dormancy once temperatures become favourable (Tiyayon, 2008). In northern growing regions, such as Ontario, Canada, unpredictable warm spells in late winter can signal catkins into blooming prematurely. Blooming catkins are notably susceptible to cold damage and premature bloom leaves catkins vulnerable to any extreme weather (-20□C) that might follow (Capik & Molnar, 2014). Cold-damaged catkins can result in significant reduction of nut yield for hazelnut farmers (Capik & Molnar, 2014). Therefore, it is of interest for the hazelnut industry in Ontario to extend the dormant state of catkins beyond any early spring-like conditions (Dale et al., 2012).

Flower dormancy is composed of three stages, including paradormancy, endodormancy and ecodormancy (Lang et al., 1987). Paradormancy is the initial cessation of growth, in response to shortening days and cold temperatures. During this stage, flowers can still bloom if placed in forcing conditions (Liu & Sherif, 2019; Yang et al., 2021). This kind of dormancy is not self-regulated, but the flower is regulated by hormones produced by other organs such as leaves (Lang et al., 1987). Endodormancy is considered a ‘true’ state of dormancy which is self-regulated and marked by the inability to resume growth even under favourable conditions (Yang et al., 2021). In ecodormancy, growth of the flower is suppressed primarily due to unfavourable environmental conditions, and can resume growth once exposed to favourable conditions (Liu & Sherif, 2019). Catkins will transition from endodormancy to ecodormancy once a threshold of time in cool temperatures (also known as the chill requirement) has been reached (Mehlenbacher, 1991). In hazelnut, the chill requirement has previously been measured for many accessions by Mehlenbacher (1991), who measured the number of chill hours required to exit endodormancy. A chill hour, in this case, was defined as an hour spent between 0-7□C (Mehlenbacher, 1991). Each accession has a unique chill requirement (Mehlenbacher, 1991).

Efforts to breed hazelnut varieties with ideal bloom times are currently underway (Valentini et al., 2021). However, breeding is a long process in hazelnut, taking almost 20 years to release an accession from its conception (Botta et al., 2019). Thus, remediation strategies to delay catkin bloom are highly desirable. The bloom of some deciduous woody perennials, such as grape and peach, has been delayed through the exogenous application of hormones (Bowen et al., 2016; Zhang & Dami, 2012; Crisosto et al., 1989; Durner & Gianfagna, 1991; Grijalva-Contreras et al., 2011; Liu & Islam et al., 2021). Indeed, plant hormones are well known to be regulators of dormant tissues, including flower dormancy, vegetative bud dormancy, axillary bud dormancy and seed dormancy (Liu & Sherif, 2019; Horvath et al., 2003; Beveridge et al., 2023; Ali et al., 2022). Thus, it is of interest to have a comprehensive understanding of the hormonal dynamics which regulate flower dormancy, so that it can be manipulated to influence bloom in deciduous woody crops with greater efficacy.

In the present study, the role of the canonical plant hormones abscisic acid (ABA), gibberellin (GA), auxin, cytokinin (CTK) and ethylene, in regulating hazelnut catkin dormancy were investigated. Usually, studies investigating the hormonal regulation of dormancy in deciduous woody perennials measure one or two hormones at a time. Additionally, it is usually only the bioactive forms of hormones that are targeted. To date, large emphasis has been put on characterizing the role of ABA and GA in dormancy regulation, perhaps due to their well-documented control of seed dormancy. It is widely observed that endogenous ABA content peaks during endodormancy and decreases during the release of dormancy. GA content generally accumulates towards release of endodormancy or ecodormancy. However, the patterns for endogenous GA content do not appear to be well conserved across studies. Auxin, cytokinin, and ethylene are vastly understudied compared to ABA and GA, with only a handful of studies measuring these hormones during dormancy. The present study measured ABA, GA, auxin, CTK and many of their metabolites, along with the ethylene precursor ACC in hazelnut catkins. By measuring all five of the hormones simultaneously, we provide greater insight into how these hormones interact in their regulation of flower dormancy. Additionally, by measuring many metabolites for each hormone, not just the bioactive forms, we observe trends in metabolic flux that would otherwise be hidden. The relevant metabolic pathways of these hormones can be seen in Figures 1–5. Hormone profiles were generated in four hazelnut accessions with varying bloom times, to determine if there were any differences in hormonal trends that could be associated with bloom phenotype. Ethylene metabolic and signaling-gene expression was also measured alongside ACC to gain greater insight into ethylene biosynthesis during dormancy.

**Figure 1.**
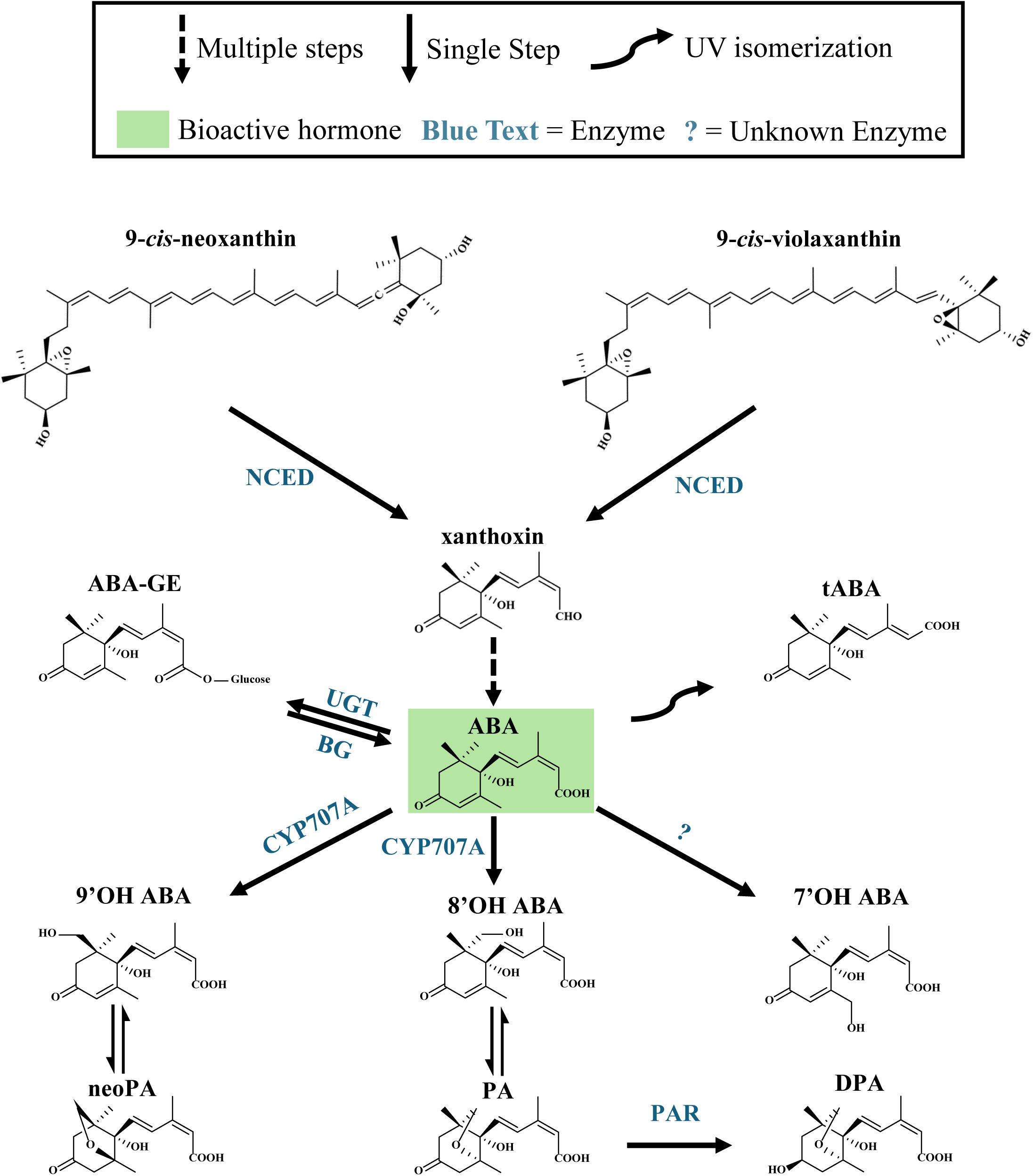
The ABA metabolic pathway. The first committed and rate-limiting step in ABA biosynthesis is the cleavage of 9′-cis-violaxanthin or 9′-cis-neoxanthin to form xanthoxin, catalyzed by 9′-cis-epoxycarotenoid dioxygenases (NCEDs) in the plastid (Seo & Marion-Poll, 2019; Wu et al., 2023). Through a series of steps, xanthoxin is converted into ABA in the cytosol (Seo & Marion-Poll, 2019; Wu et al., 2023). ABA can be spontaneously isomerized into 2-trans-ABA (tABA), which is biologically inactive, when exposed to light (Kim et al., 1992). The major catabolic route of ABA is initiated by 8′ hydroxylation in the cytosol, through the Cytochrome P450 monooxygenase (CYP) 707A to form 8′-hydroxy-ABA (8′OH ABA) (Seo & Marion-Poll, 2019; Wu et al., 2023). 8′OH ABA is unstable and spontaneously isomerizes into phaseic acid (PA) (Seo & Marion-Poll, 2019). PA is reduced to form dihydrophaseic acid (DPA) by PA reductase (PAR) (Seo & Marion-Poll, 2019). Other, minor, catabolic routes of ABA are initiated by 9′ hydroxylation and 7′ hydroxylation (Seo & Marion-Poll, 2019). 9′-hydroxy-ABA (9′OH ABA) is formed as a side-product by CYP707A (Seo & Marion-Poll, 2019). 9′OH ABA is unstable and spontaneously isomerizes into neophaseic acid (neoPA) (Seo & Marion-Poll, 2019). The enzyme for 7′ hydroxylation is currently unknown (Seo & Marion-Poll, 2019). ABA can also be inactivated via conjugation in the cytosol, with ABA glucosyl ester (ABA-GE) being the most prominent conjugate (Seo & Marion-Poll, 2019; Wu et al., 2023). ABA is conjugated by uridine diphosphate glucosyl transferases (UGT’s) (Seo & Marion-Poll, 2019). Two glucosidases, BG1 and BG2, hydrolyze ABA-GE and provide an alternate route for ABA synthesis (Lee et al., 2006; Xu et al., 2012).

**Figure 2.**
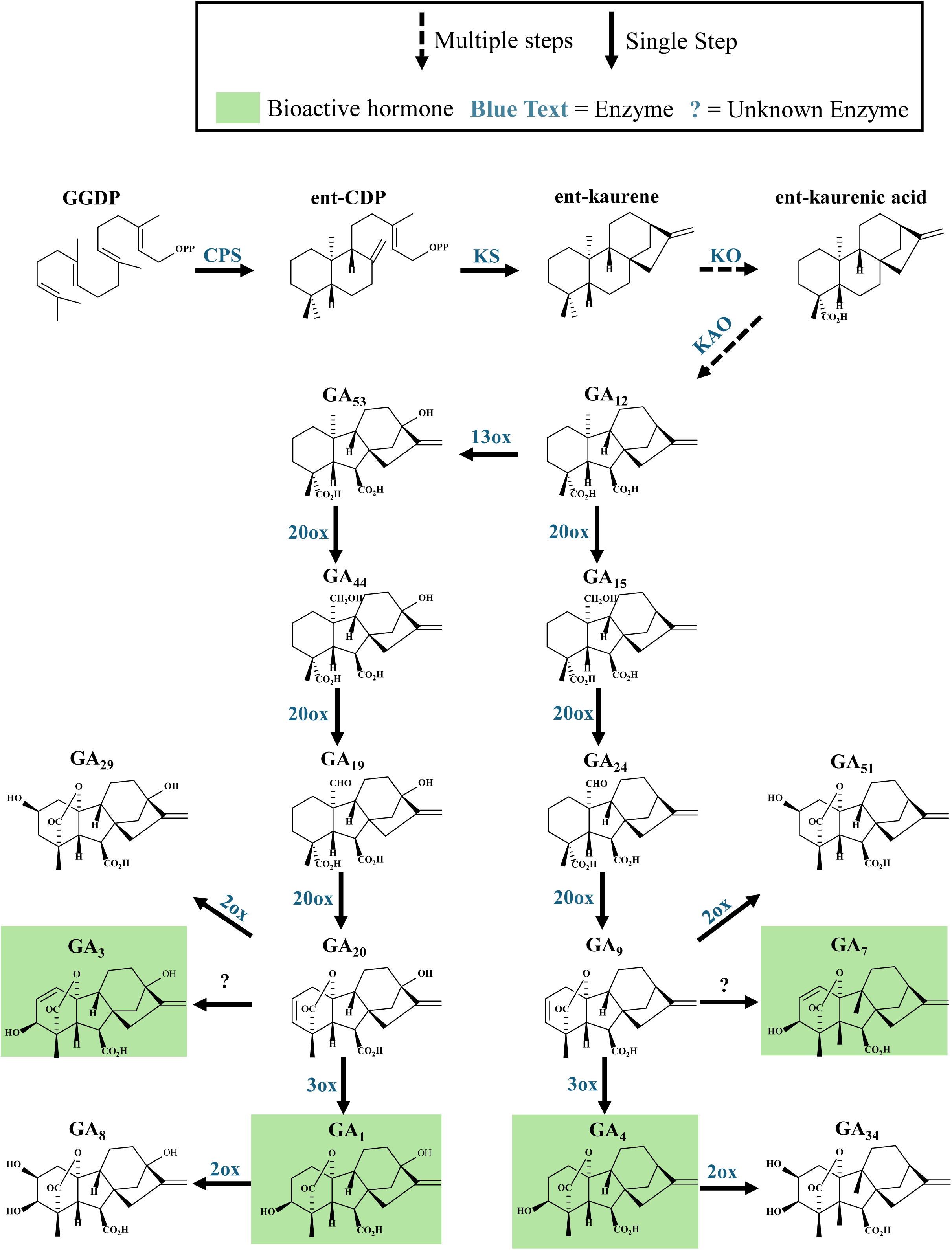
The GA metabolic pathway. The first committed step of GA biosynthesis is the conversion of geranyl geranyl diphosphate (GGDP) to ent-copalyl diphosphate (ent-CDP) by ent-copalyl diphosphate synthase (CPS) (Sun, 2008; He et al., 2020). Ent-Kaurene is subsequently formed by ent-kaurene synthase (KS) in the plastid (Sun, 2008). Ent-kaurene is oxidized in a series of reactions, facilitated by ent-kaurene oxidase (KO), which is embedded in the outer plastid membrane, to form ent-kaurenoic acid (Sun, 2008; Yamaguchi, 2008). Ent-kaurenoic acid is further oxidized by ent-kaurenoic acid oxidase (KAO) in the endoplasmic reticulum to produce GA_12_ (Sun, 2008; Yamaguchi, 2008). GA_12_ is either oxidized by 13ox to form GA_53_, to begin the GA_1_ biosynthesis pathway, or will be oxidized by 20ox to begin the GA_4_ biosynthesis pathway (Sun, 2008). In the GA_1_ biosynthesis pathway, GA_53_ oxidation is catalyzed by 20ox to produce GA_44_, GA_19_, and GA_20_, sequentially (Sun, 2008). GA_20_ is then oxidized by 3ox to form GA_1_ (Sun, 2008). In the GA_4_ pathway, GA_12_ is oxidized in a series of reactions catalyzed by 20ox to form GA_15_, GA_24_, and GA_9_, sequentially (Sun, 2008). GA_9_ is then further oxidized by 3ox to form GA_4_ (Sun, 2008). There is evidence to suggest that GA_3_ and GA_7_ are formed from the oxidation of GA_9_ and GA_20_ by 3ox, respectively, although a formal pathway has not been elucidated (Ward et al., 2010; He et al., 2020; Yamaguchi, 2008). GA is primarily deactivated by GA 2-oxidases (2ox), in the cytosol, which belong to the ODD family of enzymes and has its own small gene family (Yamaguchi, 2008). Certain 2ox oxidize bioactive GA_1_ and GA_4_, to form GA_8_ and GA_20_, respectively. GA_9_ and GA_20_, can also be oxidized by 2ox to form GA_51_, and GA_29_, respectively (Yamaguchi, 2008).

**Figure 3.**
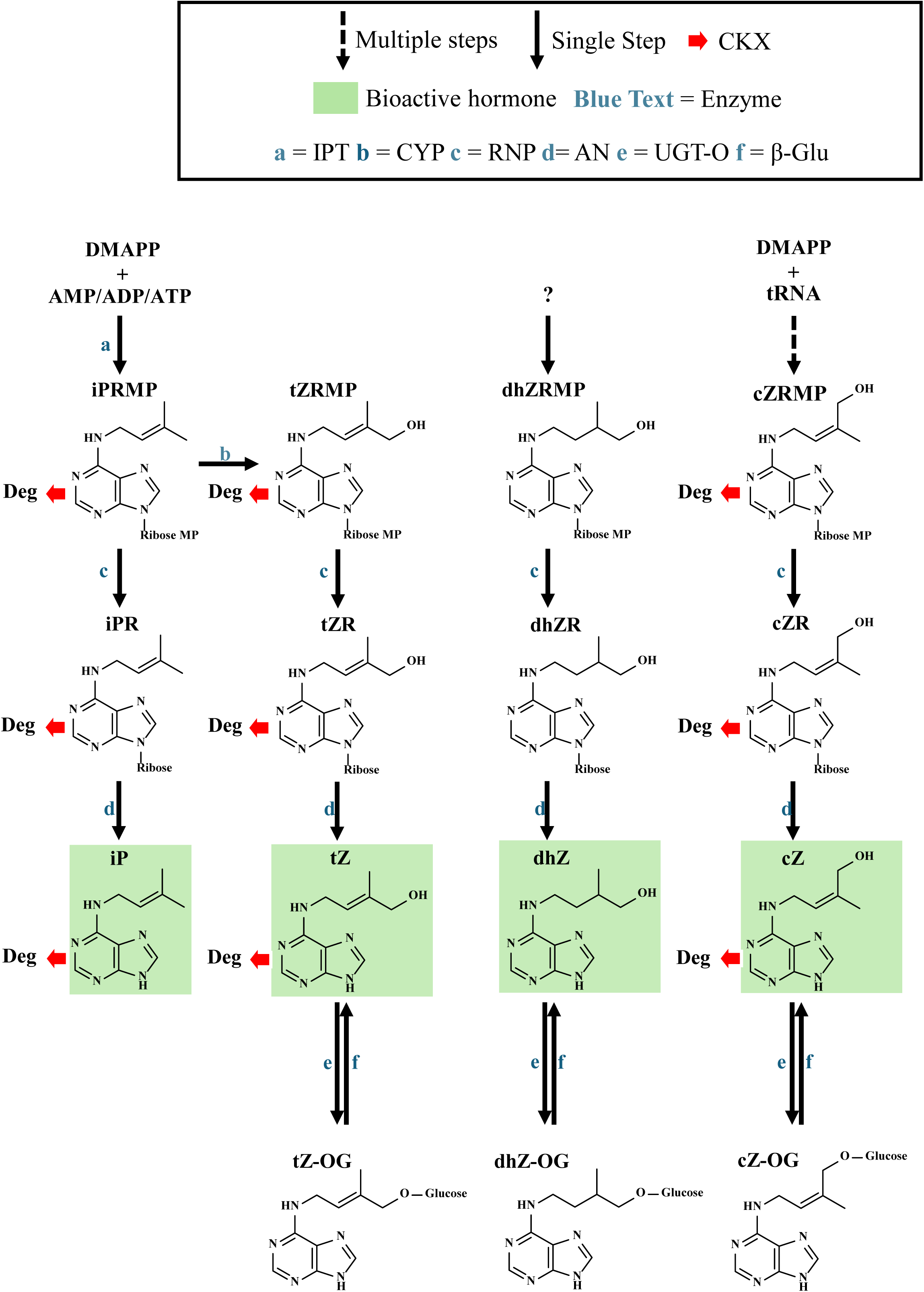
The IAA metabolic pathway. The predominant auxin in plants is indole acetic acid (IAA) (Casanova-Sáez et al., 2021). The main biosynthetic pathway, starting with tryptophan (Trp), is two steps and is believed to take place in the cytosol (Casanova-Sáez et al., 2021). Trp is first deaminated by TRYPTOPHAN AMINOTRANSFERASE OF ARABIDOPSIS 1 (TAA1) to form indole-3-pyruvic acid (IPyA) (Casanova-Sáez et al., 2021). IPyA is then decarboxylated by flavin-containing monooxygenases from the YUCCA (YUC) family to form IAA, which is the rate-limiting step (Casanova-Sáez et al., 2021). IAA can be deactivated by amino acid conjugation (Casanova-Sáez et al., 2021). The IAA-amide bond is formed by IAA acyl acid amido synthetases from the GRETCHEN HAGEN3 (GH3) family (Casanova-Sáez et al., 2021). IAA-Ala and IAA-Leu are thought to be storage conjugates which can be hydrolyzed to release free IAA by IAA-LEUCINE RESISTANT1 (ILR1) and IAA-ALANINE RESISTANT3 (IAR3), respectively (Casanova-Sáez et al., 2021). IAA-Asp and IAA-Glu are the main conjugates and can also be hydrolyzed to release free IAA by ILR1 in *A. thaliana* (Hayashi et al., 2021). The oxidation of IAA-Asp and IAA-Glu by DIOXYGENASE FOR AUXIN OXIDATION (DAO) is main, committed step of IAA deactivation in *A. thaliana* (Hayashi et al., 2021). 2-oxindole-3-acetic acid-Asp (oxIAA-Asp) and 2-oxindole-3-acetic acid-Glu (oxIAA-Glu are then hydrolyzed by ILR1 to release 2-oxindole-3-acetic acid (oxIAA) (Hayashi et al., 2021). Indole-3-butyric acid (IBA), created endogenously by the plant, is a molecule which can induce auxin activity, likely through creation of IAA, although it is unknown if it is a storage molecule or a precursor via an alternative biosynthetic pathway (Casanova-Sáez et al., 2021).

**Figure 4.**
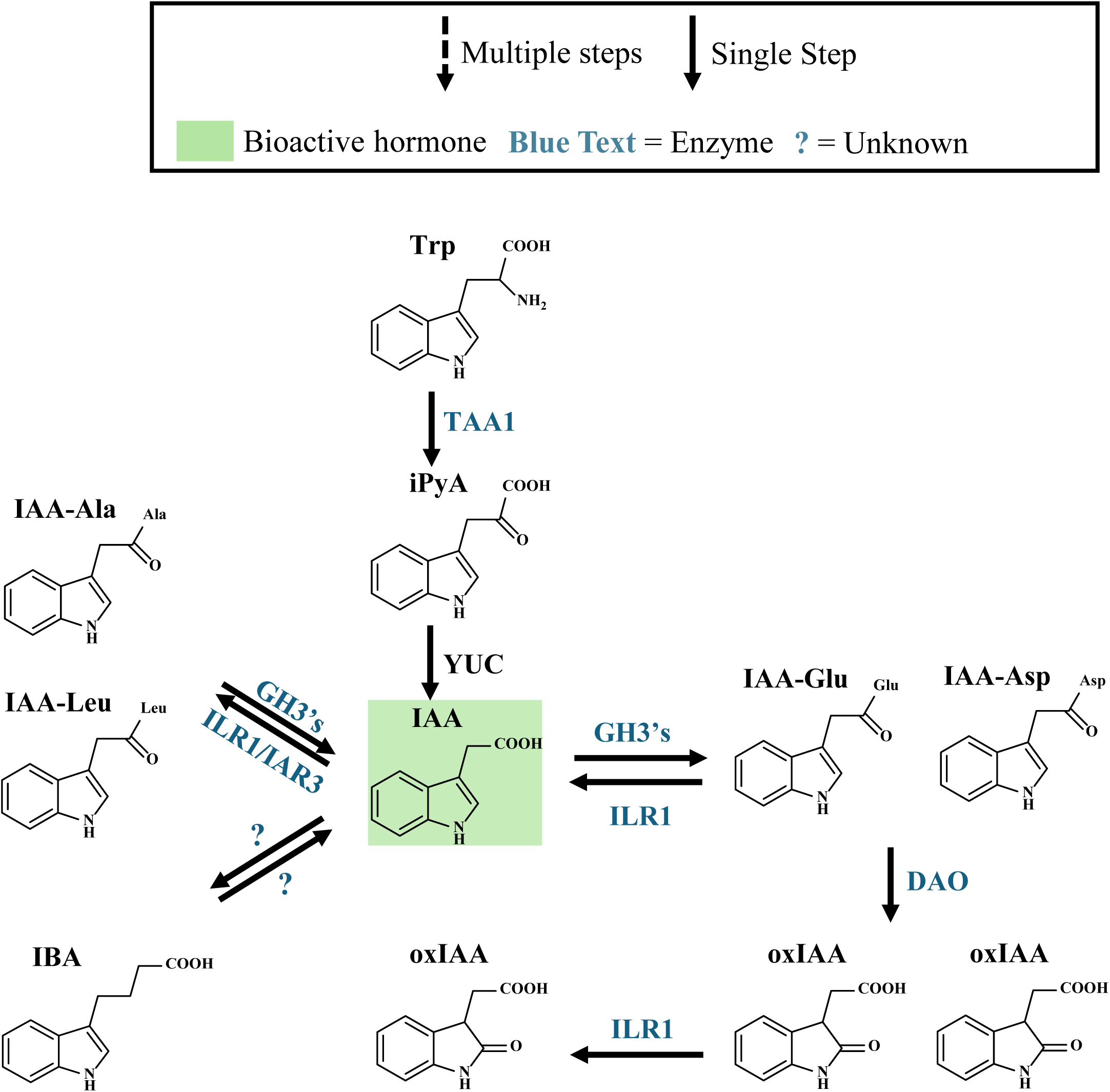
The CTK metabolic pathway. The main pathway for the biosynthesis of CTKs is the isopentenyladenosine-5′-monophosphate (iPRMP) dependent pathway (Hošek et al., 2019). The iPRMP-dependent pathway begins with the addition of the isoprenoid chain of dimethylallyl diphosphate (DMAPP) to adenosine diphosphate (ADP) or adenosine triphosphate (ATP) by adenylate isopentenyltransferase (IPT), to form isopentenyladenosine-5′-diphosphate (iPRDP) or isopentenyladenosine-5′-triphosphate (iPRTP), respectively, which are dephosphorylated to form iPMRP in the plastid (Hošek et al., 2019; Zhao et al., 2024). iPRMP can be hydroxylated to form tZ riboside monophosphate (tZRMP) in the endoplasmic reticulum (Hošek et al., 2019; Zhao et al., 2024). iPRMP and tZRMP are converted to iP riboside (iPR) and tZ riboside (tZR) by a 5′-ribonucleotide phosphohydrolase (RNP), respectively, in the cytosol (Hošek et al., 2019; Zhao et al., 2024). dhZ riboside monophosphate (dhZRMP) is similarly converted to dhZ, although the origin of dhZRMP is unknown (Hošek et al., 2019). cZ riboside monophosphate (cZRMP), derived from tRNAs, is converted to cZ riboside (cZR) and cZ in the cytosol (Hošek et al., 2019; Zhao et al., 2024). CTK ribosides are converted to free bases by adenosine nucleosidases (ANs) in the cytosol or apoplast (Hošek et al., 2019; Zhao et al., 2024). iP, tZ, and cZ are primarily deactivated by the irreversible cleavage of their side chain, via CTK oxidase/dehydrogenase (CKX) (Hošek et al., 2019). Deactivation by CKX could occur in the cytosol, apoplast, vacuole, endoplasmic reticulum or the nucleus (Zhao et al., 2024). tZ, dhZ, and cZ can be reversibly glucosylated at the O position by zeatin-O-glucosyltransferase (UGT-O) to deactivate CTKs (Hošek et al., 2019). trans-zeatin-O-glucoside (tZ-OG), dihydrozeatin-O-glucoside (dhZ-OG), and cis-zeatin-O-glucoside (cZ-OG) can be reactivated by β-glucosidase (β-GLU) (Hošek et al., 2019). Glucosylation at the O position is suspected to take place in the cytosol and the chloroplast (Zhao et al., 2024).

**Figure 5.**
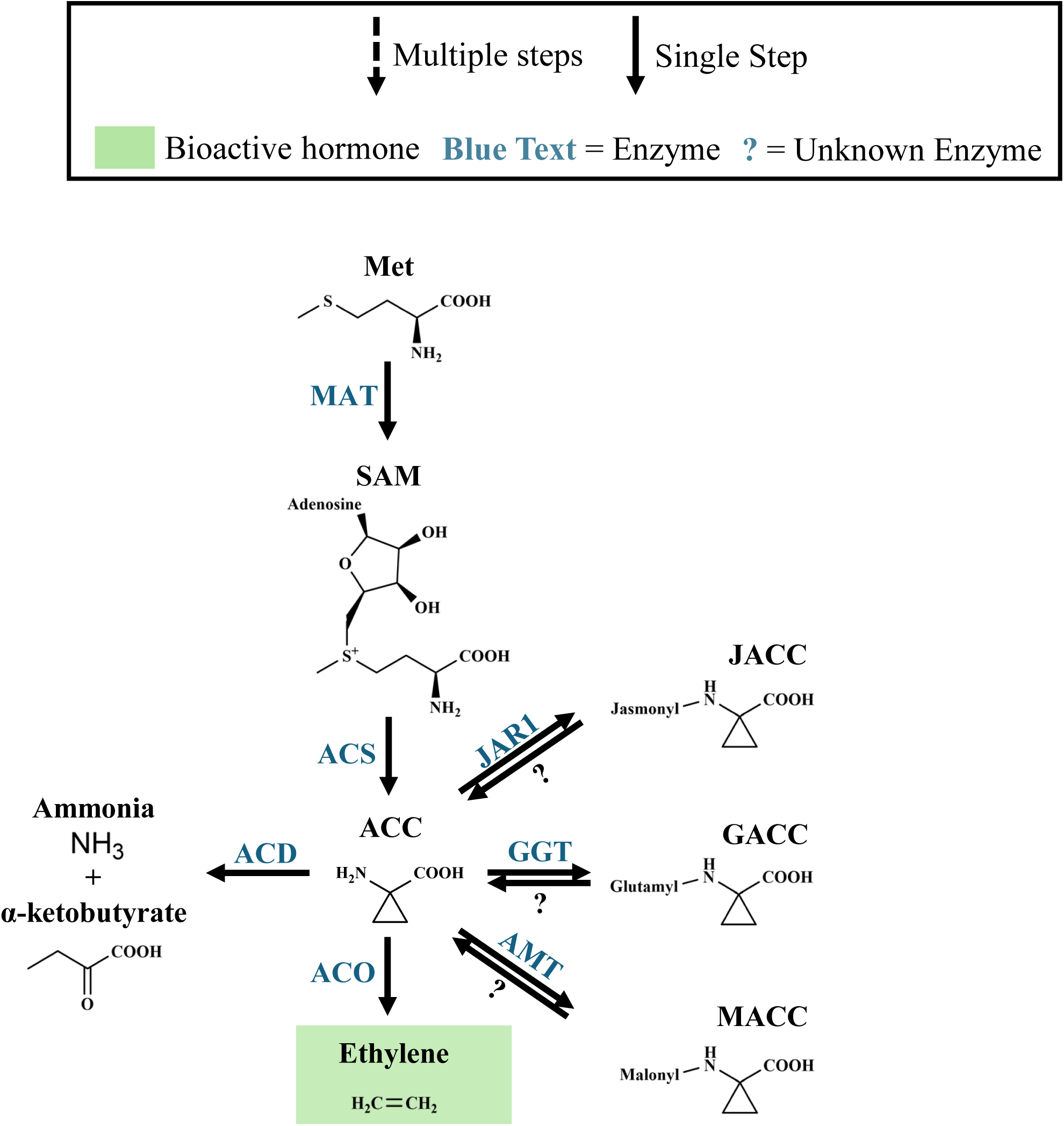
The ethylene metabolic pathway. Ethylene biosynthesis begins with the creation of S-adenosyl methionine (SAM) from methionine (Pattyn et al., 2021). SAM is then converted into 1-aminocyclopropane-1-carboxylic acid (ACC) by ACC synthase (ACS) in the cytosol (Pattyn et al., 2021; Kahn et al., 2024). ACC is oxidized by ACC oxidase (ACO) to produce ethylene (Pattyn et al., 2021). The subcellular localization of ACO is uncertain, with evidence suggesting either the plasma membrane or the cytosol (Kahn et al., 2024). ACC can be conjugated to form malonyl-ACC (MACC), glutamyl-ACC (GACC), and jasmonyl-ACC (JACC) by ACC-N-malonyl transferase (AMT), γ□glutamyl transpeptidase (GGT), and Jasmonic Acid Resistance 1 (JAR1), respectively (Pattyn et al., 2021). MACC, GACC, and JACC can be reconverted into ACC by unknown enzymes. ACC can also be broken down into α-ketobutyrate and ammonia by ACC deaminase (ACD) (Singh et al., 2015).

## Materials & Methods

### Sample collection

Hazelnut trees were planted at the University of Guelph’s Simcoe Research Station in 2009 and 2010. During the 2021/2022 season, multiple studies were carried out simultaneously by other research groups at the Research Station. *C. avellana* x *C. colourna* ‘Farris G-17’ #’s 182, 183, 184, and *C. avellana* ‘Jefferson’ #’s 239, 240, 241, as well as an open-pollinated *C. americana* ‘Native’ #318 tree were included in the study. All trees were grown in the south field of the hazelnut research plot and accessions were within 50 m from each other. Trees of the same label were directly adjacent to one another. Catkins were collected approximately every two weeks on September 7^th^, and 27^th^, October 12^th^, and 25^th^, November 8^th^, and 22^nd^, December 6^th^ and 20^th^ of 2021 as well as January 4^th^, 18^th^ and 31^st^, February 14^th^, and 28^th^, March 10^th^, and 25^th^ of 2022. Due to logistical concerns surrounding travel, collections took place in the late morning to early afternoon (10 a.m. to 1 p.m.) from October 12^th^, 2021 to January 4^th^, 2022, January 31^st^, 2022, and February 28^th^, 2022. Collections took place early afternoon to late afternoon (1 p.m. to 5 p.m.) on September 7^th^ and 27^th^, 2021 as well as January 18^th^, February 14^th^, March 10^th^ and 25^th^, 2022.

On each collection date, approximately five catkins per tree were randomly selected and had their length measured with calipers while still attached to the tree. Then, 12 catkins per tree were randomly selected from around the tree, placed in 4.5 mL, internally threaded, cryogenic storage vials (Thermo Fisher Scientific, #12-567-502) and flash-frozen in liquid nitrogen within approximately 2 min of being picked. Three catkins were allocated for hormone profiling, gene expression analysis, pollen development analysis, and as backups, respectively. Frozen catkins were transported to Queen’s University using a 4.1 L, Cryo Vapor Shipper (CX100B-11M, Taylor-Wharton, NW Bolivar, Ohio) and finally stored between -70 to -80□C in a cryogenic freezer.

Some of the trees were being treated with different fertilizers or mulch as a part of a separate study performed by researchers at the Simcoe Research Station. ‘Jefferson’ #239 received 20:20:20 Nitrogen:Phosphorous:Potassium (N:P:K) plus extra K, ‘Jefferson’ #241 received 20:20:20 N:P:K plus extra P, ‘Farris’ #182 received K plus mulch, ‘Farris’ G-17 #183 received N plus mulch, ‘Farris’ G-17 #184 received Magnesium (Mg) plus extra micronutrients, while ‘Jefferson’ #240 and the *C. americana* tree received no treatments.

### Hormone profiling

Three whole catkins per tree were lyophilized. While handling catkins prior to lyophilization, catkins were held in liquid nitrogen or in a Corning® CoolRack® M90 Module (Corning, Cat # 432044) on dry ice to remain cryogenic. The CoolRack®, containing the catkins, was transferred directly from dry ice into a SP VirTis Advantage EL-85 Benchtop Shelf Freeze Dryer (ATS Scientific Products, Warminster, Pennsylvania), which took 2 min to reach 1000 millitorr and reached a final pressure of approximately 50 millitorr. Catkins were left to lyophilize with the condenser set to -55°C and the shelf set to 20°C. After 48 hours, catkins were weighed with an analytical balance accurate to 0.1 mg, and were weighed again after 4 to 6 hours, or overnight, until the weight decreased less than 1 mg from the previous measurement. Lyophilized catkins were ground in a dry mortar & pestle. Per tree, approximately 20 mg of ground tissue from each catkin was pooled. Pooled, ground tissue was sent to the National Research Council in Saskatoon for total hormone profiling.

Thirty-eight metabolites were targeted including: ABA, ABA-GE, DPA, PA, 7′OH-ABA, neo-PA, tABA, GA_1_, GA_3_, GA_4_, GA_7_, GA_8_, GA_9_, GA_19_, GA_20_, GA_24_, GA_29_, GA_34_, GA_44_, GA_51_, GA_53_, IAA, IBA, IAA-Asp, IAA-Glu, IAA-Ala, IAA-Leu, cZ, tZ, iP, dhZ, cZR, tZR, cZOG, tZOG, dhZR, iPR, and ACC. Analysis was performed on a UPLC/ESI-MS/MS. The procedure for quantification of ABA and ABA catabolites, CTKs, auxins, and gibberellins in the hazelnut tissue samples was performed using a modified procedure described in Lulsdorf et al. (2013), while quantification of ACC was performed using a modified procedure described in Chauvaux et al. (1997). These modifications have not been published.

### Identification of *C. avellana* ethylene metabolic and signaling gene homologs

*C. avellana ACC SYNTHASE* (*ACS*), *ACC OXIDASE* (*ACO*), *ACC DEAMINASE* (*ACD*) and ethylene receptor genes were initially identified by querying the *C. avellana* ‘Tombul’ genome (NCBI RefSeq assembly GCF_901000735.1) with the Basic Local Alignment Search Tool (BLAST) on the NCBI website. Protein sequences of *A. thaliana* homologs were used as queries for tBLASTn (Table 1). Because the *C. avellana* ‘Tombul’ genome was only resolved at the chromosome level on NCBI at the time of study, a putative homolog was identified if it had approximately 50% sequence identity to the query sequence and the majority of the query was spread across supposed exons in close proximity to each other (i.e. supposed introns between sequential, supposed exons were of a reasonable size). For each putative gene, a query (or queries) with the highest query coverage and percent identity was used to localize presumptive exons. Exons were then pieced together manually to create a predicted amino acid sequence for each putative homolog. Putative homologs had their identity confirmed by using predicted amino acid sequences as a query against *A. thaliana* sequences using BLASTp (protein BLAST) on NCBI. If the top, annotated BLAST hit was indeed a true *A. thaliana* homolog, the putative *C. avellana* homolog was retained. The predicted coding sequences of *C. avellana* genes were used to design qPCR primers.

**Table 1.**
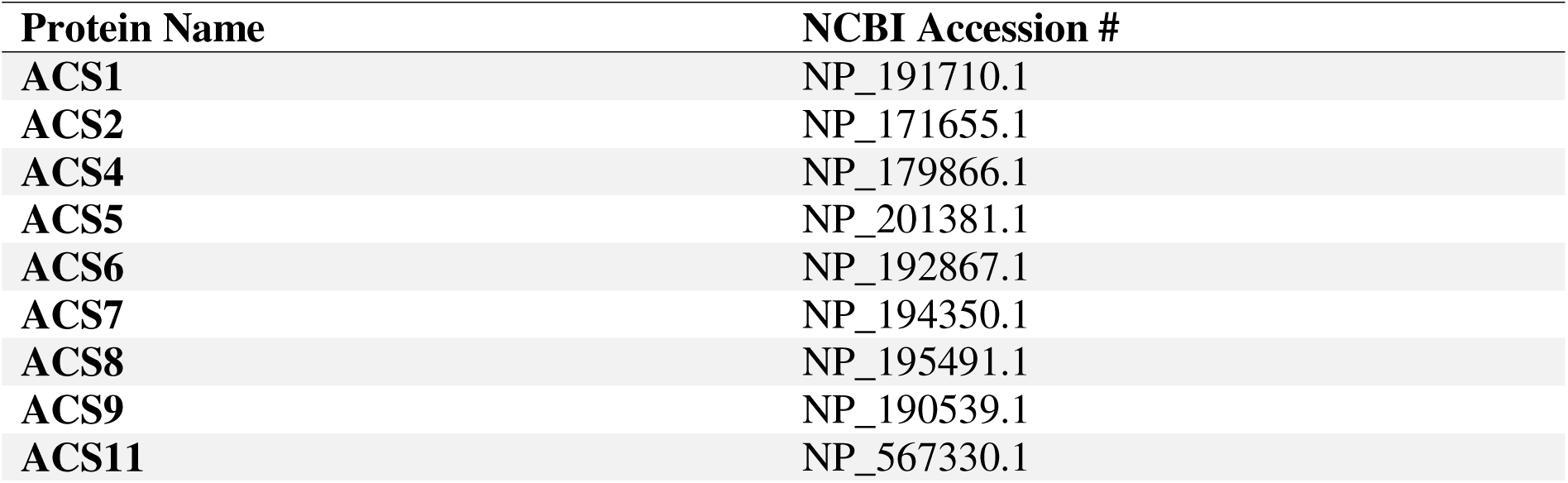
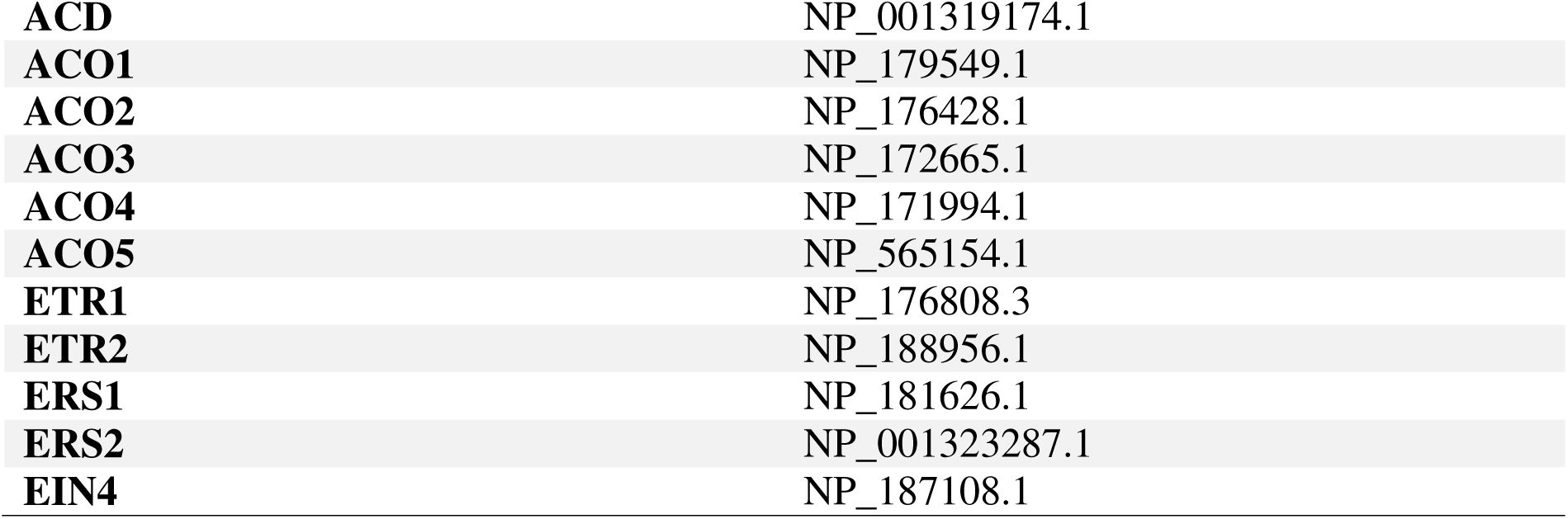
Listed are the names and NCBI accession #’s of *A. thaliana* ACS, ACD, ACO, and ethylene receptor proteins used as queries for tBLASTn and BLASTp on NCBI.

| Protein Name | NCBI Accession # |
| --- | --- |
| ACS1 | NP_191710.1 |
| ACS2 | NP_171655.1 |
| ACS4 | NP_179866.1 |
| ACS5 | NP_201381.1 |
| ACS6 | NP_192867.1 |
| ACS7 | NP_194350.1 |
| ACS8 | NP_195491.1 |
| ACS9 | NP_190539.1 |
| ACS11 | NP_567330.1 |
| **ACD** | NP_001319174.1 |
| **ACO1** | NP_179549.1 |
| **ACO2** | NP_176428.1 |
| **ACO3** | NP_172665.1 |
| **ACO4** | NP_171994.1 |
| **ACO5** | NP_565154.1 |
| **ETR1** | NP_176808.3 |
| **ETR2** | NP_188956.1 |
| **ERS1** | NP_181626.1 |
| **ERS2** | NP_001323287.1 |
| **EIN4** | NP_187108.1 |

### Phylogenetic Analysis

Once available on NCBI, the gene annotations and official predicted protein sequences of *C. avellana* ‘Tombul’ were used to complete a phylogenetic analysis. The *A. thaliana* and *C. avellana* homologs, listed in Tables 1 and 2, respectively, for ACS, ACO and ethylene receptor proteins were used to generate a multiple sequence alignment using the MUSCLE algorithm. These multiple sequence alignments were used for subsequent analysis of the evolutionary history of *A. thaliana* and *C. avellana* ACS, ACO and ethylene receptor proteins. The evolutionary history of proteins was inferred using the Maximum Likelihood method, with the Le_Gascuel_2008 model for ACS and ACO and the JTT matrix-based model for ethylene receptors (Le & Gascuel, 2008; Jones et al., 1992). A discrete Gamma distribution was used to model evolutionary rate differences among sites. Phylogenetic analysis of *C. avellana* ACS, ACO and ethylene receptor proteins was completed using MEGA11: Molecular Evolutionary Genetics Analysis version 11 (Tamura et al., 2021).

**Table 2.** Listed are the names and NCBI accession #’s of *Corylus avellan*a ACS, ACO, and ethylene receptor proteins used for phylogenetic analysis in MEGA11.

| **Protein Name** | **NCBI Accession #** |
| --- | --- |
| **ACS2** | XP_059439399.1 |
| **ACS3** | XP_059442427.1 |
| **ACS8** | XP_059462029.1 |
| **ACS11** | XP_059431684.1 |
| **ACO2** | XP_059437717.1 |
| **ACO5** | XP_059449964.1 |
| **ACO6** | XP_059455434.1 |
| **ACO11** | XP_059432141.1 |
| **ETR1** | XP_059440948.1 |
| **ETR2** | XP_059447186.1 |
| **EIN4** | XP_059437359.1 |

### DNA extraction

DNA was extracted from hazelnut catkins using the DNeasy® Plant Pro Kit (Qiagen, #69204), Approximately 100 mg pieces of whole catkins were used as starting material for the DNeasy® Plant Pro Kit. Manufacturer protocols were followed. DNA quality and quantity was estimated using the DeNovix DS-11+ Spectrophotometer (DeNovix Inc., Wilmington, Delaware) microvolume analysis.

### RNA extraction

RNA was extracted using the Norgen Biotek Plant/Fungi Total RNA Purification Kit (Norgen Biotek, # 25800). Catkins were removed from the -80□C freezer and placed in a hand-held dewar containing liquid nitrogen to keep them at cryogenic temperatures while handling. A mortar and pestle were chilled by adding liquid nitrogen, until approximately 75% full, and liquid nitrogen was left to evaporate. This was done a total of three times to ensure complete chilling of the mortar and pestle. A whole catkin was added to the chilled mortar and pestle, while liquid nitrogen was present, and ground into a fine powder. Liquid nitrogen was re-added shortly after evaporation to ensure tissue integrity. Grinding was complete after three to four evaporation cycles. Approximately 25-50 mg of tissue was added to 600µL of lysis buffer C + 6µL 2-mercaptoethanol (Sigma-Aldrich, M3148) and mixed by vortexing. Samples were incubated at 55□C for five min, mixing 2 to 3 times during incubation via inversion. Sample was added to a filter column and spun for 2 min at 20,000 x *g*. Clear supernatant was transferred to a 1.5 mL microcentrifuge tube and an equal volume of 100% ethanol was added. Samples were mixed by vortexing. Half of the sample was added to a binding column and centrifuged for 1.5 min at 5000 x *g*. This step was repeated for the remaining sample. Wash solution A, 400 µL, was added to the column and centrifuged for 1.5 min at 20,000 x *g*. This wash step was repeated two more times. The column was centrifuged for 2.5 min at 20,000 x *g* to dry the column. Columns were taken out of the centrifuge and left on the benchtop for two min with the lid open to allow for ethanol evaporation. The column was centrifuged for 2.5 min at 20,000 x *g* once more. Elution buffer, 50 µL, was added to the column and left to incubate for two min. RNA was eluted by spinning at 20,000 x *g* for 1.5 min.

RNA was electrophoresed on a 1% agarose, 1% bleach, TAE (40 mM Tris base, 20 mM acetic acid, 1 mM Ethylenediaminetetraacetic acid) gel to observe RNA integrity (Aranda et al., 2012). RNA was retained if a 25S rRNA:18S rRNA ratio of over 1 was observed. The 25S rRNA:18S rRNA ratio was determined with densitometry using imageJ software (Schneider et al., 2012). RNA quality and quantity was estimated using the DeNovix DS-11+ Spectrophotometer microvolume analysis. RNA with concentrations above 100 ng/µL had 260/230 ratios of approximately 1.8 and 260/280 ratios of approximately 2, indicating acceptable purity. RNA with concentrations below 100 ng/µL had 260/230 ratios of approximately 1.5 and 260/280 ratios of approximately 1.8. A low 260/230 and 260/280 ratio for these samples was primarily ascribed to lower nucleic acid content, rather than increased presence of contaminants from catkin tissue, and was therefore deemed acceptable. RNA was treated with DNAse using the Invitrogen Turbo DNA-free™ kit (Thermo Fisher Scientific, Cat # AM1907). Up to 5000 ng of RNA was added to a 25µL DNAse reaction containing 1µL of Turbo DNAse (2 Units/µL). The reaction was incubated at 37°C for 25 min. The reaction was halted by adding 5 µL of inactivation reagent. DNAse-treated RNA once again had its RNA quantity estimated using the DeNovix DS-11+ Spectrophotometer microvolume analysis. RNA was stored in a -80□C freezer until further use.

### cDNA preparation

Complementary DNA (cDNA) was prepared using the LunaScript® RT SuperMix Kit (New England Biolabs, Cat #E3010S). RNA, 500 ng, was combined with 5 µL of 4X Supermix or 4X - RT mix along with provided water to prepare 20 µL cDNA and no reverse transcriptase (-RT) control reaction mixes, respectively. cDNA reaction mixes and -RT controls had a final RNA concentration of 25 ng/µL. Reaction mixes were incubated at 25□C for 2 min, 55□C for 10 min and 95□C for 1 min in a T100™ Thermal Cycler (Bio-Rad, # 1861096). cDNA was kept frozen at -20°C until needed.

### qPCR

qPCR reactions contained 2 µL of 1/12.5 diluted cDNA, 5 µL of 2x PowerTrack™ SYBR Green Master Mix, 0.25 µL 40X Yellow Sample Buffer included with the 2X PowerTrack™ SYBR Green Master Mix, appropriate amounts of 10 µM forward and reverse primers each, and molecular biology grade water to a final volume of 10 µL. Three technical replicates were included per sample. A no template (molecular biology grade water) and a positive control (1/100 dilution of 35 ng/µL ‘Jefferson’ gDNA, extracted using the DNeasy® Plant Pro Kit) was included for each target on every plate. A -RT control was done, with three technical reps, for each sample at least once, for the chaActin qPCR assay. Reactions were loaded into 96-well, thin-wall Bio-Rad Hard-Shell® PCR Plates (Bio-Rad, Cat #HSP9601) and sealed with an adhesive, optical Microseal ‘B’ PCR Plate Sealing Film (Bio-Rad, Cat #MSB1001). Reaction master mix and sample were mixed by spinning the plate. qPCR was run on the Bio-Rad CFX96TM Real-Time System + C1000 TouchTM Thermal Cycler (Bio-Rad Laboratories Inc., Mississauga, Canada). Reactions were initially denatured at 95□C for 3 min. Amplification cycles included a 95□C denaturation step for 5 sec and an annealing/elongation step at the respective TM for 30 sec. A total of 40 amplification cycles were completed. A melt curve was completed after each run to check for non-specific amplification. Not all samples/timepoint or all timepoints/sample could be included on one plate. Therefore, the chaActin qPCR assay, using a 1/100 dilution of 35 ng/µL ‘Jefferson’ gDNA as a template, was included in triplicate as an inter-plate calibrator to account for inter-plate variability. Each target was run at its ideal TM, by using a gradient TM setting on each run. Each target was also run using its ideal primer concentration, tested between 200 – 400 nM in 50 nM increments. Primer efficiency was determined according to the formula 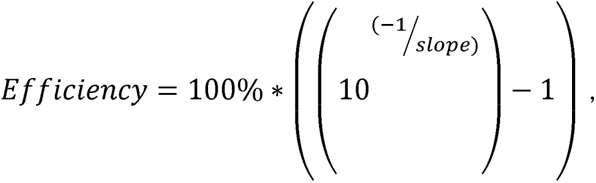 where “slope” is the slope of a Log_10_ base efficiency curve with 1/10, 1/20, 1/40, 1/80 and 1/160 serial dilutions of various template gDNA or cDNA. A list of primers for each target can be seen in Table 3.

**Table 3.**
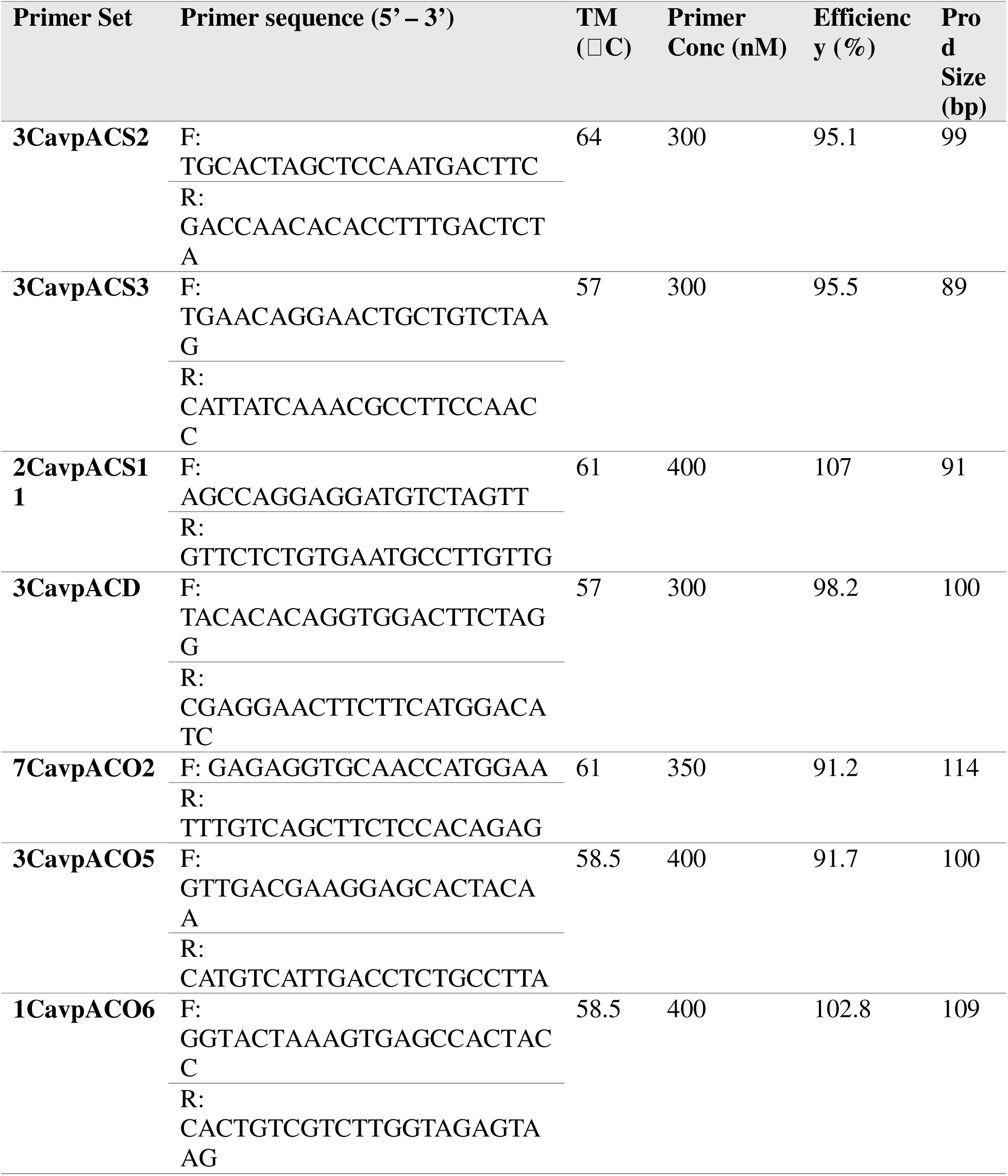
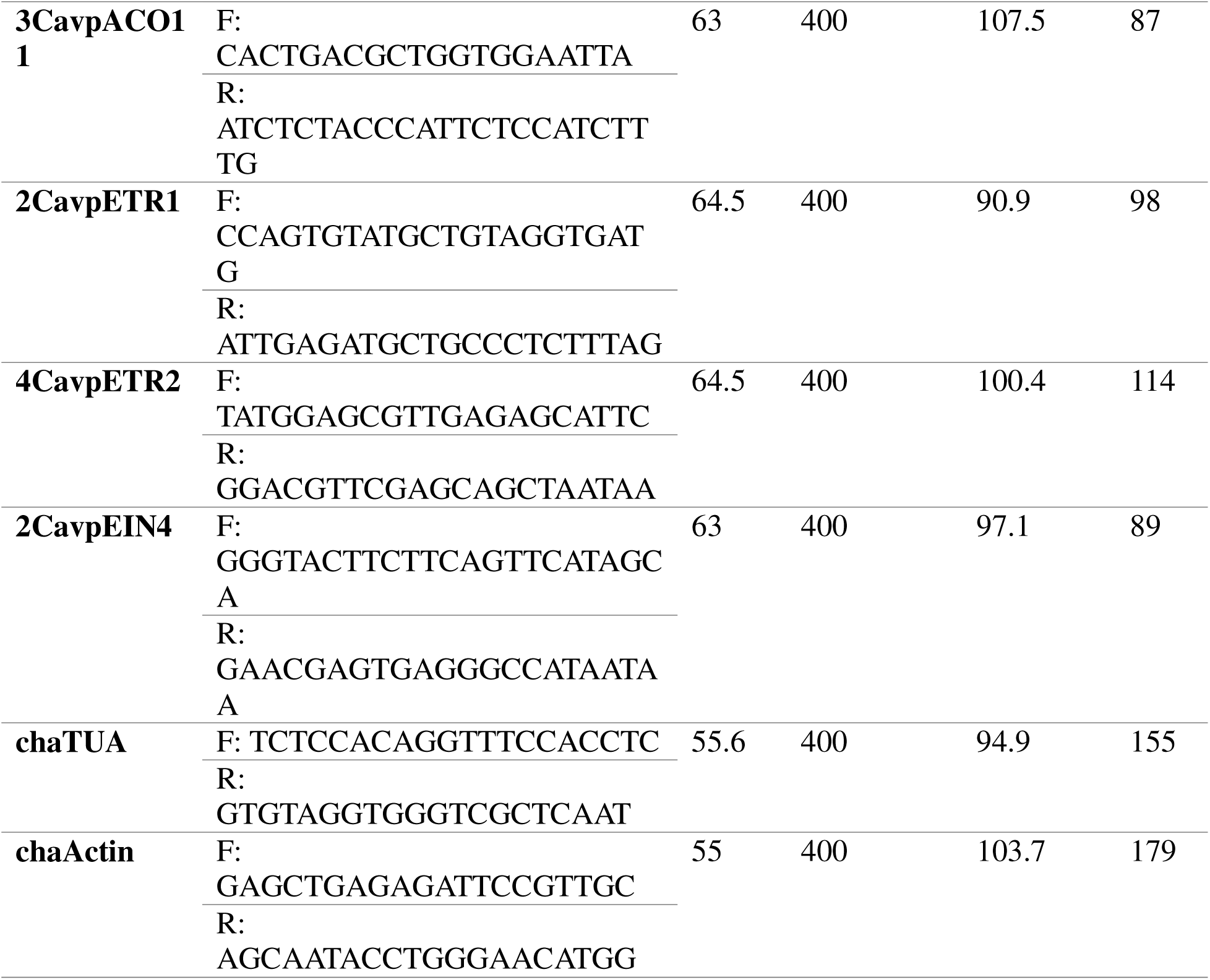
qPCR Primer Sets for *C. avellana* Metabolic and Receptor Genes. Listed are the primer sets used to measure the relative transcript abundance of *C. avellana ACS2, ACS3, ACS11, ACD1, ACD2, ACO2, ACO5, ACO2, ACO5, ACO6, ACO11, ETR1, ETR2, EIN4, TUA*, and *ACTIN* genes, along with their optimized TM, primer concentration, and efficiency. chaTUA and chaActin primers were obtained from Hou et al. (2021).

To determine the relative expression of each target, the Cq values of each sample were first corrected for inter-plate variability. This was done by subtracting the inter-plate calibrator Cq value (from the plate on which the target was run) from the target Cq value, then adding the average inter-plate calibrator Cq value (the average from across every qPCR plate run). The corrected Cq values for each target were used for subsequent determination of relative expression. Relative expression was determined for a given target and sample according to the following formula: : 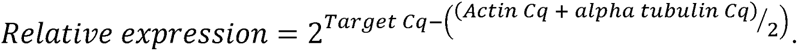

## Results

### Weather and catkin phenotype during the 2021/2022 season

Catkins were collected from three hazelnut accessions with different expected bloom dates. *C. avellana* x *C. colourna* ‘Farris G-17’ (n=2), *C. avellana* ‘Jefferson’ (n=3), and a wild *C. americana* (n=1) have early, early, mid, and late blooming phenotypes, respectively. ‘Farris G-17′ typically blooms about three weeks earlier than ‘Jefferson’, although the magnitude of difference is highly dependent on the weather of a given season. Initially, a third ‘Farris G-17′ (‘Farris G-17’ #182) was included in the study. As the season progressed, however, this tree appeared to have shorter catkins with greater girth and different colouration compared to the other two ‘Farris G-17′ trees. This tree, relabelled ‘Farris 182’, was suspected to be a different accession, mislabeled upon planting, and was analyzed separately from ‘Farris G-17′. The lineage of ‘Farris 182’ is currently unknown. Although ‘Farris 182’ is likely genetically distinct from ‘Farris-G17′, observations made during the 2022/2023 season suggest that it has a similar early blooming phenotype. Thus, ‘Farris 182’ was included in the analysis as a single tree with an early-bloom phenotype.

Catkins were collected bi-weekly from September 2021 to March 2022. Photographs of catkins and trees were taken upon each collection date (Supplementary material). By November 8th, most of the leaves had begun turning, and the trees were bare by December 6^th^. Catkin lengths were monitored on collection dates to estimate dormancy establishment in the fall and the beginning of bloom in the spring (Figure 6A). Catkins appear to have stopped elongating by September 27th for ‘Farris 182’ and by October 12th for all other accessions (Figure 6A). Since it was a cold winter, all of the accessions initiated bloom within the same two-week period between March 10th and March 25^th^ (Figure 6A). Photographs of the catkins show ‘Farris 182’ and ‘Farris G-17′ are more advanced in bloom stage compared to ‘Jefferson’, and ‘Jefferson’ is more advanced in bloom stage than *C. americana* (Figure 6B). Although most catkins did not initiate bloom until mid-late march, ‘Farris G-17′ had a handful of catkins which appeared to bloom pre-maturely mid-January (supplementary material). The bulk of chill hours was accumulated by catkins between mid-October and late December (Figure 6C).

**Figure 6.**
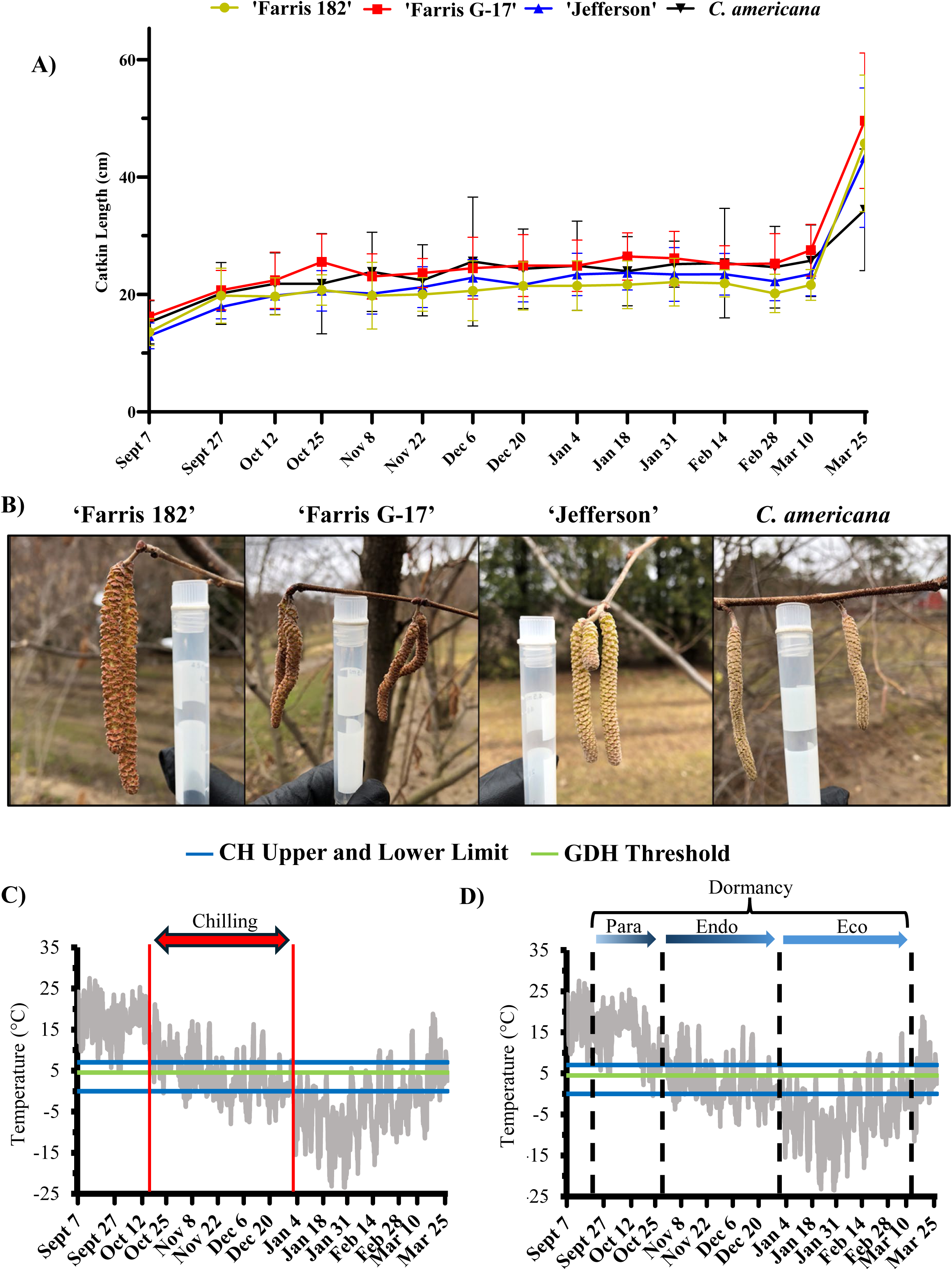
Catkin phenotype and environmental temperature during the 2021/2022 season. **A)** Shown are the average catkin lengths (mm) from September 7th, 2021 to March 25th, 2022. Error bars represent standard deviation. ‘Farris 182’ catkins appear to have reached full length by September 27^th^, while the other accessions achieved full catkin length by Oct 12^th^. All catkins began elongating between March 10^th^ and March 25^th^. **B)** Representative images of hazelnut catkins photographed March 25^th^, 2022. A Fisherbrand™ internally threaded, 4.5 mL cryogenic storage vial (Cat no. 12-567-502) is included for scale. ‘Farris 182’ and ‘Farris G-17′ catkins are fully open and shedding pollen. ‘Jefferson’ catkins are partially open, shedding a small amount of pollen. *C. americana* catkins are just starting to open. **C)** The average hourly temperature retrieved from the Environment Canada, Delhi, Ontario station, approximately 25 km from the Simcoe Research Station. The upper and lower limits of chill hour (CH) accumulation as used by Mehlenbacher (1991) as well as the threshold for growing degree hour (GDH) accumulation as reported by Tiyayon (2008) are included. Red vertical bars outline the period in which the majority of chill hours were accumulated. **D)** The average hourly temperature retrieved from the Environment Canada, Delhi, Ontario station, approximately 25 km from the Simcoe Research Station. The upper and lower limits of chill hour (CH) accumulation as used by Mehlenbacher (1991) as well as the threshold for growing degree hour (GDH) accumulation as reported by Tiyayon (2008) are included. Catkins were likely paradormant on September 27th and October 12th; endodormant between October 25th and January 4th; ecodormant between January 4th and March 10th. Vertical dashed lines indicate likely dormancy transitions. Colour gradients within arrow bars reflect the gradual transitions between dormancy states likely to occur.

### Assigning dormancy states based on temperature data

A traditional dormancy curve could not be completed, and no reliable biomarker to track dormancy in hazelnut catkins has been discovered. Therefore, the dormancy status of catkins was estimated using catkin length and temperature data. Catkins had halted growth by September 27^th^ and were likely paradormant at this stage (Figure 6D). The transition to endodormancy is difficult to determine visually, but it is likely that it occurred during October, as most of the trees leaves began desiccating by November. The transition from endodormancy to ecodormancy likely took place during November and December, while the majority of CH were being accumulated (Figure 6D). Catkins were likely fully ecodormant for the remainder of the study, after which few CH were accumulated (Figure 6D). It is likely that transitions between para and endodormancy, as well as endo and ecodormancy, were gradual (Figure 6D). i.e. on Oct 12th the catkins were likely transitioning to endodormancy, and on November 8th the catkins were likely in a deeper stage of endodormancy than on December 6th. The dormant status of the catkins, as discussed here, is superimposed on relevant figures going forward. Hormone profiling was completed for eight collection dates throughout the season. Two dates during paradormancy (Sept 27^th^ and Oct 12^th^), two dates in endodormancy (Nov 8^th^ and Dec 6^th^), and four dates in ecodormancy (Jan 18^th^, Feb 14^th^, Feb 28^th^ and Mar 10^th^) were sent for hormone profiling.

### Hormone Profiling of Hazelnut Catkins

#### ABA

Of the ABA metabolites measured, ABA, ABA-GE, DPA, and PA were the predominant metabolites found in the catkins whereas 7′OH-ABA, neo-PA, and tABA were considered negligible. The data for the three negligible metabolites can be seen in the supplementary material. Of the abundant metabolites, DPA was predominant throughout the season (Figure 7 A,B,C,D). Generally, ABA had higher levels in the fall than in the winter, with levels peaking during endodormancy (Figure 7E). In ecodormancy, ABA decreased and remained low, leading toward bloom (Figure 7E). PA displayed a similar trend (Figure 7 A,B,C,D). In contrast to ABA and DPA, ABA-GE levels were low in the onset of dormancy, increased during endodormancy, and remained high in ecodormancy leading toward bloom (Figure 7 A,B,C,D). Despite large differences in flux through the ABA biosynthetic pathway, represented by DPA accumulation, all accessions had a similar amount of bioactive ABA throughout the season (Figure 7F).

**Figure 7.**
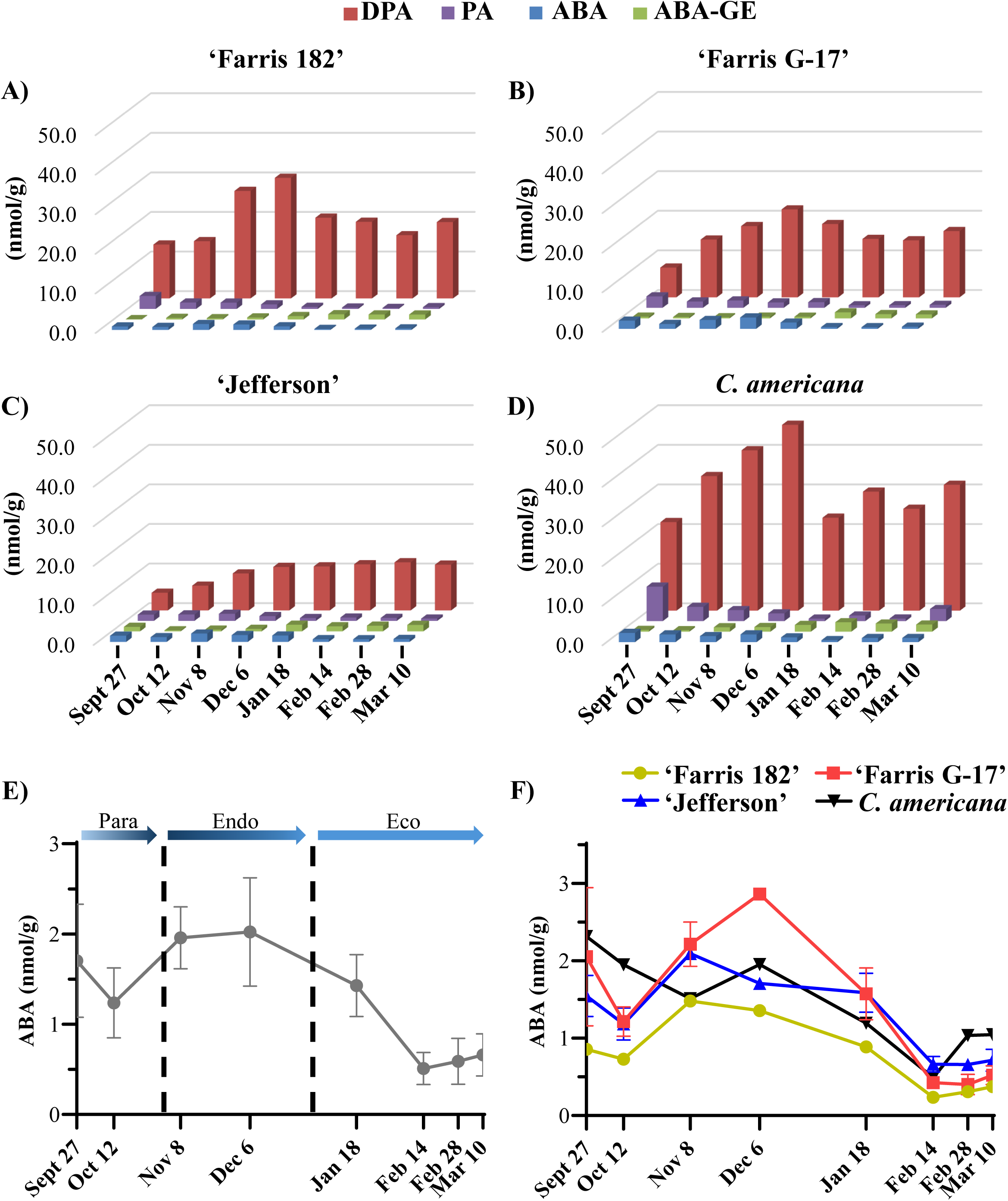
ABA metabolites measured in catkins during the 2021/2022 season. A) The concentration (nmol/g) of ABA, ABA-GE, PA, and DPA in ‘Farris 182’ catkins (n=1). (B) The concentration (nmol/g) of ABA, ABA-GE, PA, and DPA in ‘Farris G-17’ catkins (n=2). (C) The concentration (nmol/g) of ABA, ABA-GE, PA, and DPA in ‘Jefferson’ catkins (n=3). (D) The concentration (nmol/g) of ABA, ABA-GE, PA, and DPA in *C. americana* catkins (n=1). E) The average concentration (nmol/g) of ABA in catkins in all accessions. Error bars indicate standard deviation. Vertical dashed lines indicate likely dormancy transitions. Colour gradients within arrow bars reflect the gradual transitions between dormancy states likely to occur. “para” = paradormancy, “endo” = endodormancy, “eco” = ecodormancy. F) The concentration (nmol/g) of ABA in in ‘Farris 182’ (n=1), ‘Farris G-17′ (n=2), ‘Jefferson’ (n=3), and *C. americana* (n=1) catkins. Error bars represent standard deviation.

#### 2.3.2 GA

Of all the GAs measured in this study, only GA_8_, GA_19_, GA_44_, and GA_53_ could be measured. Although none of the active GA’s (GA_1_, GA_3_, GA_4_, GA_7_) were quantifiable, it can be assumed that GA_1_ is the predominant active gibberellin in hazelnut catkins, because of the abundance of its precursors GA_19_, GA_44_, and GA_53_, and its immediate catabolite GA_8_. GA_19_ was the predominant metabolite overall (Figure 8A,B,C,D). Total GA content was pooled to visualize flux through the GA biosynthetic pathway (Figure 8E). Total GA content agrees with the trends of GA_19_, with levels low at the beginning of dormancy, gradually increasing during endodormancy, and plateauing in ecodormancy, remaining high approaching bloom (Figure 8E). GA content appears to be highest in the ‘Jefferson’ and ‘Farris 182’ accessions and lowest in the *C. americana* tree throughout the whole season (Figure 8F).

**Figure 8.**
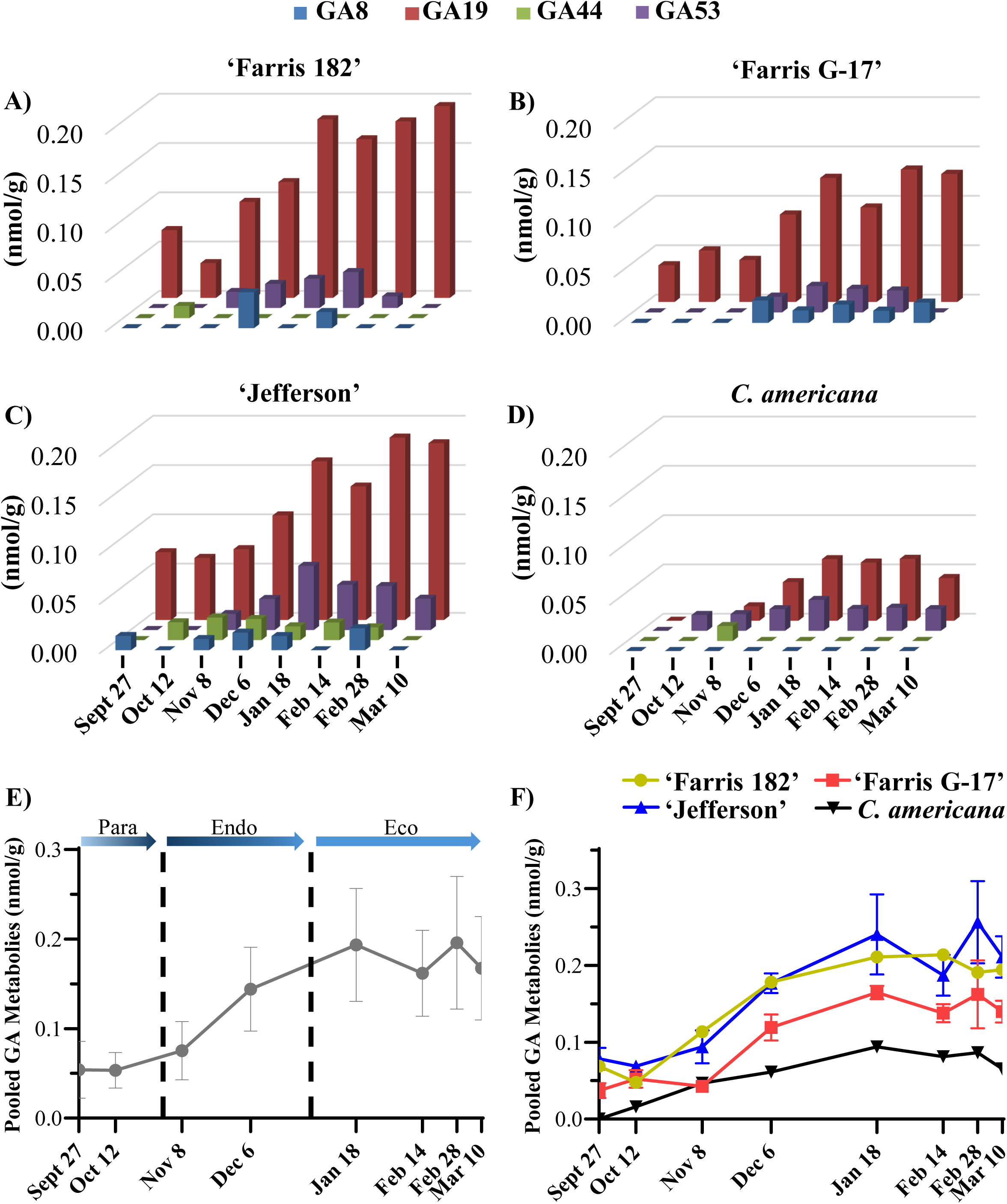
GA metabolites in catkins during the 2021/2022 season. A) The concentration (nmol/g) of GA_8_, GA_44_, GA_53_, GA_19_ in ‘Farris 182’ catkins (n=1). (B) The concentration (nmol/g) of GA_8_, GA_44_, GA_53_, GA_19_ in ‘Farris G-17’ catkins (n=2). (C) The concentration (nmol/g) of GA_8_, GA_44_, GA_53_, GA_19_ in ‘Jefferson’ catkins (n=3). (D) The concentration (nmol/g) of GA_8_, GA_44_, GA_53_, GA_19_ in *C. americana* catkins (n=1). E) The average concentration (nmol/g) of pooled GA metabolites (GA_8_, GA_44_, GA_53_, GA_19_) in catkins from all accessions. Error bars indicate standard deviation. Vertical dashed lines indicate likely dormancy transitions. Colour gradients within arrow bars reflect the gradual transitions between dormancy states likely to occur. “para” = paradormancy, “endo” = endodormancy, “eco” = ecodormancy. F) The concentration (nmol/g) of pooled GA metabolites (GA_8_, GA_44_, GA_53_, GA_19_) in in ‘Farris 182’ (n=1), ‘Farris G-17′ (n=2), ‘Jefferson’ (n=3), and *C. americana* (n=1) catkins. Error bars represent standard deviation.

#### 2.3.3 Auxin

Of the auxin metabolites measured, only IAA and IAA-Asp were detected. IAA and IAA-Asp were present in roughly equivalent amounts and shared a similar trend for ‘Jefferson” and *C. americana*, but not in ‘Farris 182’ and ‘Farris G-17’ (Figure 9A,B,C,D). IAA-Asp was largely absent in ‘Farris 182’ and ‘Farris G-17’ during endodormancy and ecodormancy (Figure 9A,B). Generally, IAA levels were high at the beginning of paradormancy and decreased toward the onset of winter, then were stable during ecodormancy (Figure 9E). IAA spiked in ‘Farris 182’, ‘Jefferson’, and *C.americana*, during ecodormancy, although the spike occurred on a different date for each accession and no such increase was observed in ‘Farris G-17’ (Figure 9F).

**Figure 9.**
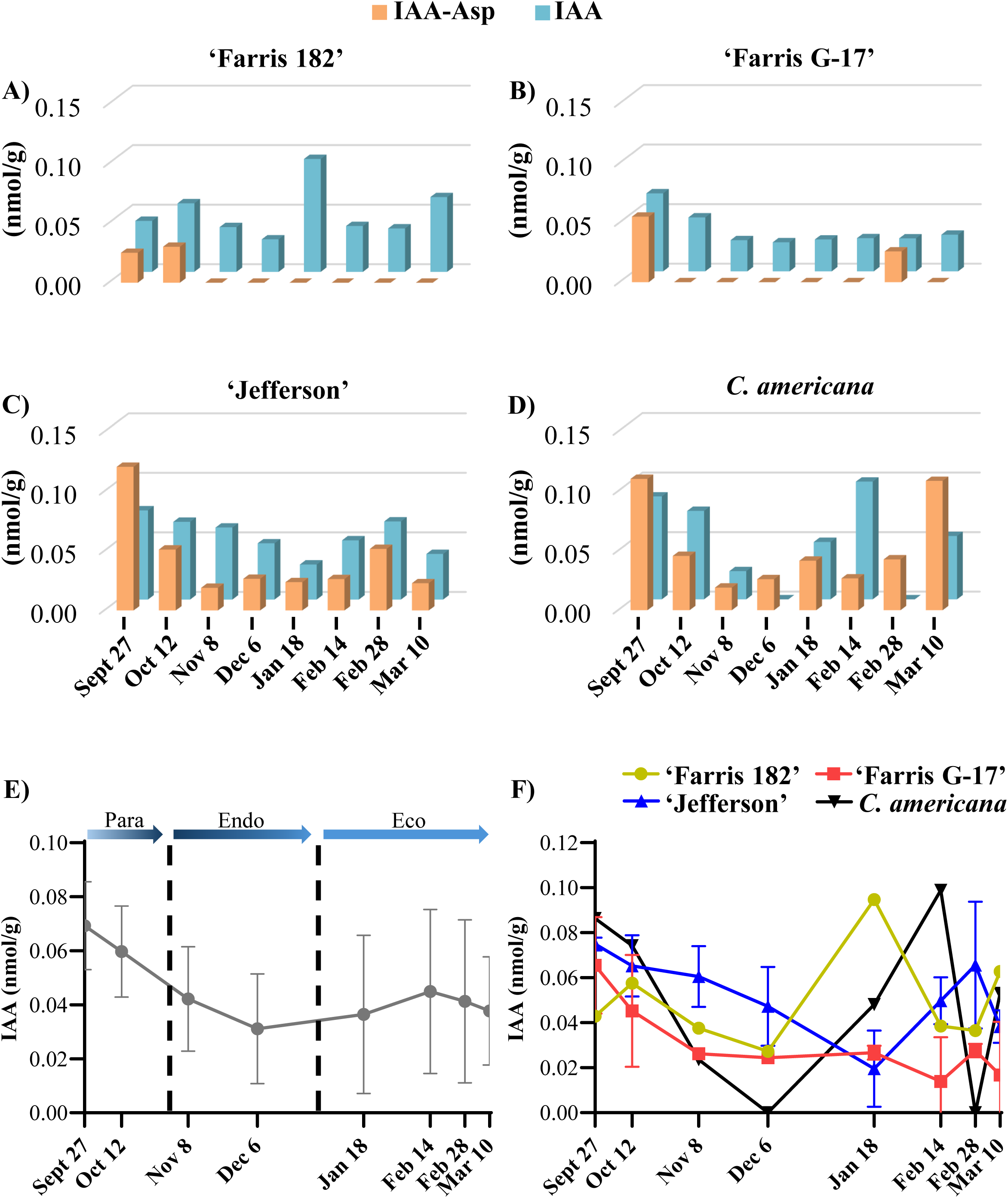
IAA and IAA-Asp in catkins during the 2021/2022 season. A) The concentration (nmol/g) of IAA and IAA-Asp in ‘Farris 182’ catkins (n=1). (B) The concentration (nmol/g) of IAA and IAA-Asp in ‘Farris G-17’ catkins (n=2). (C) The concentration (nmol/g) of IAA and IAA-Asp in ‘Jefferson’ catkins (n=3). (D) The concentration (nmol/g) of IAA and IAA-Asp in *C. americana* catkins (n=1). E) The average concentration (nmol/g) of IAA in catkins from all accessions. Error bars indicate standard deviation. Vertical dashed lines indicate likely dormancy transitions. Colour gradients within arrow bars reflect the gradual transitions between dormancy states likely to occur. “para” = paradormancy, “endo” = endodormancy, “eco” = ecodormancy. F) The concentration (nmol/g) of IAA in in ‘Farris 182’ (n=1), ‘Farris G-17′ (n=2), ‘Jefferson’ (n=3), and *C. americana* (n=1) catkins. Error bars represent standard deviation.

#### 2.3.4 CTK

Of the CTK metabolites measured, all except cZ were detected. Generally, CTK ribosides were present in much greater quantities compared to other metabolites (Figure 10, Figure 11A,B,C,D). In each accession, iP is the primary active cytokinin on Sept 27, but decreases on Oct 12^th^ and only begins to increase again after November 8^th^ (Figure 11A,B,C,D). Additionally, dhZ appears to increase drastically from Oct 12^th^ to Nov 8^th^ to quickly become the predominant active CTK in most instances (Figure 11A,B,C,D). Despite dhZ being the predominant active CTK, iPR is the primary metabolite in ‘Farris 182’ and ‘Farris G-17’ while tZR is the primary metabolite in ‘Jefferson’ and *C. americana* after December 6^th^ (Figure 10A,B,C,D).

**Figure 10.**
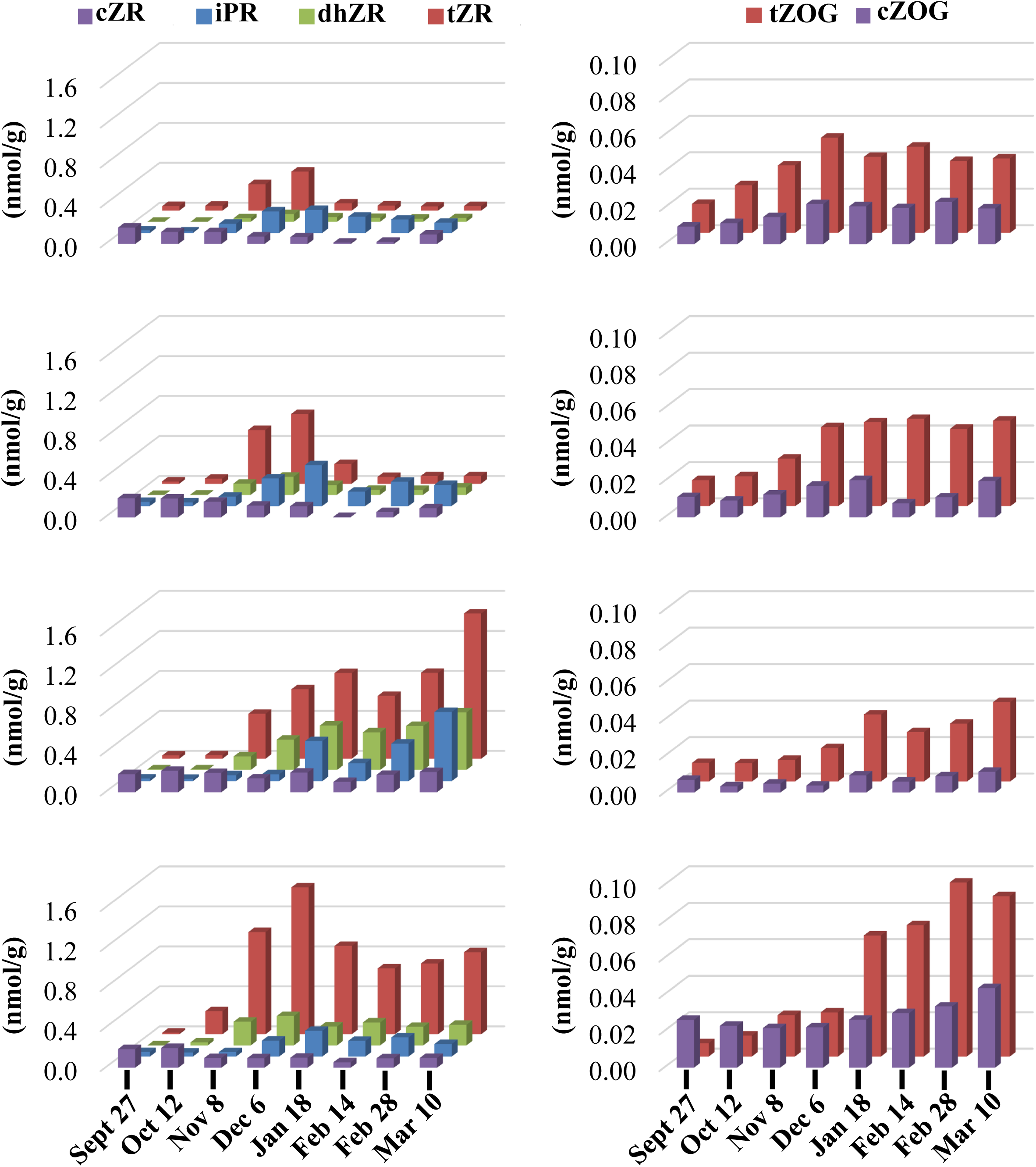
CTK ribosides and conjugates in catkins during the 2021/2022 season. A) The concentration (nmol/g) of iPR, dhZR, and tZR in ‘Farris 182’ catkins (n=1). B) The concentration (nmol/g) of cZOG and tZOG in ‘Farris 182’ catkins (n=1). C) The concentration (nmol/g) of iPR, dhZR, and tZR in ‘Farris G-17’ catkins (n=2). D) The concentration (nmol/g) of cZOG and tZOG in ‘Farris G-17’ catkins (n=2). E) The concentration (nmol/g) of iPR, dhZR, and tZR in ‘Jefferson’ catkins (n=3). F) The concentration (nmol/g) of cZOG and tZOG in ‘Jefferson’ catkins (n=3). G) The concentration (nmol/g) of iPR, dhZR, and tZR in *C. americana* catkins (n=1). H) The concentration (nmol/g) of cZOG and tZOG in *C. americana* catkins (n=1).

**Figure 11.**
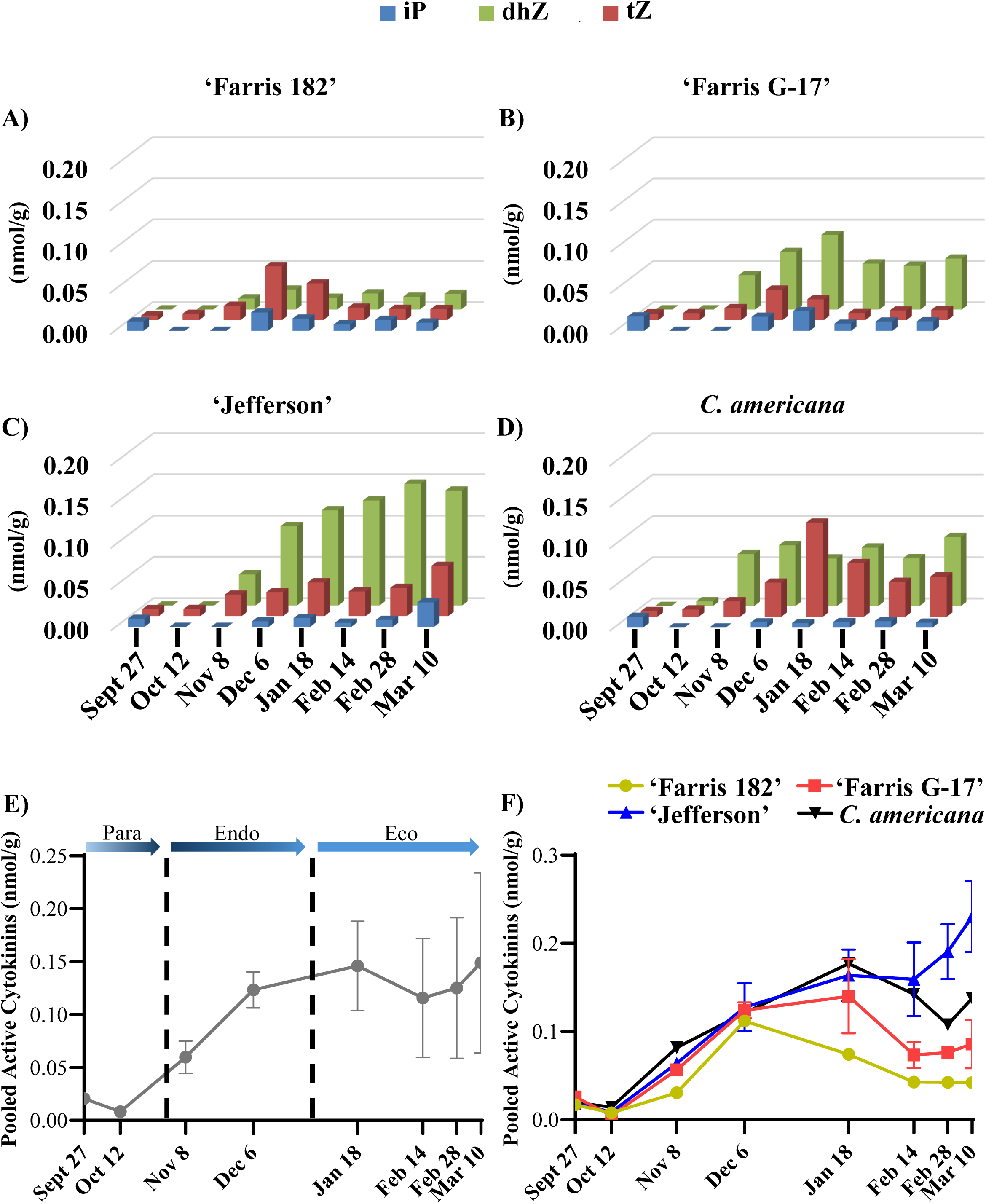
Active CTK metabolites in catkins during the 2021/2022 season A) The concentration (nmol/g) of iP, tZ, and dhZ in ‘Farris 182’ catkins (n=1). (B) The concentration (nmol/g) of iP, tZ, and dhZ in ‘Farris G-17’ catkins (n=2). (C) The concentration (nmol/g) of iP, tZ, and dhZ in ‘Jefferson’ catkins (n=3). (D) The concentration (nmol/g) of iP, tZ, and dhZ in *C. americana* catkins (n=1). E) The average concentration (nmol/g) of pooled active CTK metabolites (iP, tZ, and dhZ) in catkins from all accessions. Error bars indicate standard deviation. Vertical dashed lines indicate likely dormancy transitions. Colour gradients within arrow bars reflect the gradual transitions between dormancy states likely to occur. “para” = paradormancy, “endo” = endodormancy, “eco” = ecodormancy. F) The concentration (nmol/g) of pooled active CTK metabolites (iP, tZ, and dhZ) in in ‘Farris 182’ (n=1), ‘Farris G-17′ (n=2), ‘Jefferson’ (n=3), and *C. americana* (n=1) catkins. Error bars represent standard deviation.

Active CTKs (tZ, iP, and dhZ) were pooled to reflect total active CTK content (Figure 11E,F). Generally, active CTK content was low at the beginning of dormancy, steadily increased during endodormancy, and plateaued in ecodormancy, remaining high leading toward bloom (Figure 11E). During para and endodormancy all accessions share a similar active CTK pattern, but trends deviate between accessions in ecodormancy (Figure 11F). ‘Farris 182’ CTK levels begin to decrease in January, and plateau in February (Figure 11F). ‘Farris G-17′ and *C. americana* CTK content increases slightly in January, decreases sharply afterwards and plateaus in February (Figure 11F). ‘Jefferson’ CTK content, in contrast, peaks in January, remains high, and begins to increase further in mid-February toward bloom (Figure 11F).

#### 2.3.5 Ethylene

Generally, ACC levels were high at the beginning of paradormancy and decreased throughout endo and ecodormancy (Figure 12A). The levels of ACC varied significantly among the accessions (Figure 12B). The concentration of ACC appears to be much higher in ‘Farris 182’ and ‘Farris G-17′ compared to ‘Jefferson’ and *C. americana* accessions throughout the whole season (Figure 12B). Indeed, one-tailed, independent samples t-tests, with Welch’s correction for unequal variances, confirm ‘Farris G-17′ has greater ACC levels than ‘Jefferson’ on September 27^th^ (p=0.028), October 12^th^ (p=0.026), November 8^th^ (p=0.011), December 6^th^ (p=0.011), February 14^th^ (p=0.04) and February 28^th^ (p=0.007). There was no significant difference in ACC levels of ‘Farris G-17′ and ‘Jefferson’ on January 18^th^ (p=0.110) or March 10^th^ (p=0.054).

**Figure 12.**
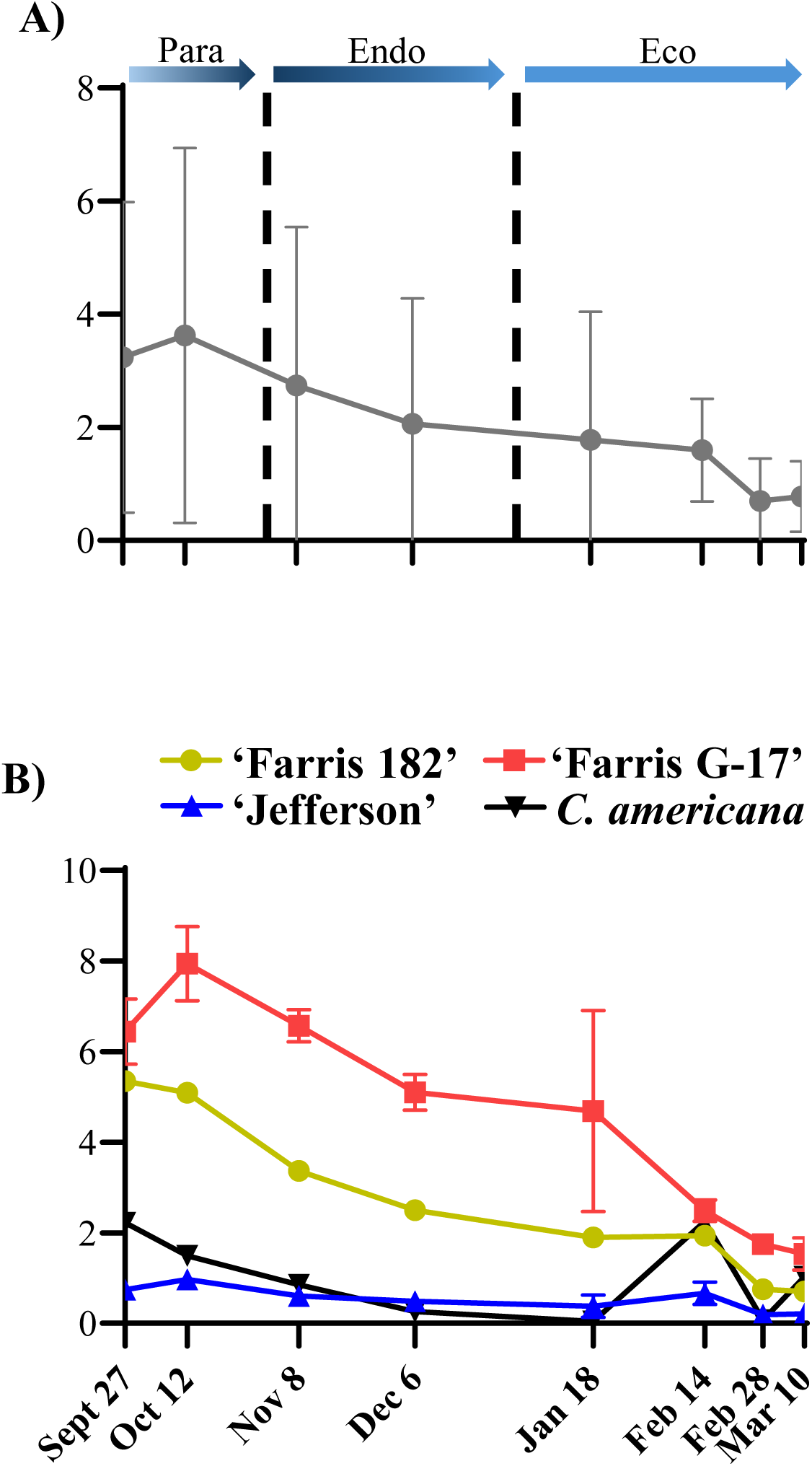
ACC in catkins during the 2021/2022 season. The average concentration (nmol/g) of ACC in catkins from all accessions. Error bars indicate standard deviation. Vertical dashed lines indicate likely dormancy transitions. Colour gradients within arrow bars reflect the gradual transitions between dormancy states likely to occur. “para” = paradormancy, “endo” = endodormancy, “eco” = ecodormancy. F) The concentration (nmol/g) of ACC in ‘Farris 182’ (n=1), ‘Farris G-17′ (n=2), ‘Jefferson’ (n=3), and *C. americana* (n=1) catkins. Error bars represent standard deviation.

### 2.4 Ethylene metabolic and signaling gene transcript abundance

#### 2.4.1 Identification of *C. avellana ACS*, *ACO*, *ACD*, and ethylene receptor genes

To gain insight as to whether ACC levels are reflective of flux through the ethylene biosynthetic pathway or are accumulating without a concomitant increase in ethylene or ethylene signaling activity, the relative transcript abundance of ethylene biosynthetic genes *ACS* and *ACO*, *ACD*, and ethylene receptors were compared to ACC levels in ‘Farris G-17′ and ‘Jefferson’.

Putative *C. avellana ACS*, *ACO*, *ACD*, and ethylene receptor genes were identified based upon homology with known *A. thaliana* genes. Four *ACS* genes (*CaACS2*, *CaACS3*, *CaACS8*, *CaACS11*), four ACO genes (*CaACO2*, *CaACO5*, *CaACO6*, *CaACO11*), three ethylene receptor genes (*CaETR1*, *CaETR2*, *CaEIN4*), and two ACD genes (*CaACD1*, *CaACD2*) were found in the *C. avellana* genome. *ACS* and *ACO* genes were labelled based upon the chromosome on which they reside, whereas ethylene receptor genes were labelled according to their homology with *A. thaliana* homologs and *ACD* genes were labelled according to the order of discovery.

#### 2.4.2 qPCR

Relative transcript abundance of *ACS*, *ACO*, *ACD*, and ethylene receptor genes was measured in catkins via qPCR for October 12^th^, December 6^th^, February 14^th^ and March 10^th^ during the 2021/2022 season. *ACS8* was not detected in any of the four timepoints, in either ‘Jefferson’ or ‘Farris G-17’ (Supplementary material). Also, *ACD1* and *ACD2* DNA sequences were nearly identical (Supplementary material) and thus a single qPCR primer set was designed to detect both transcripts. Total *ACS*, *ACO*, *ACD*, and ethylene receptor relative transcript abundances were plotted alongside ACC levels (Figure 13).

**Figure 13.**
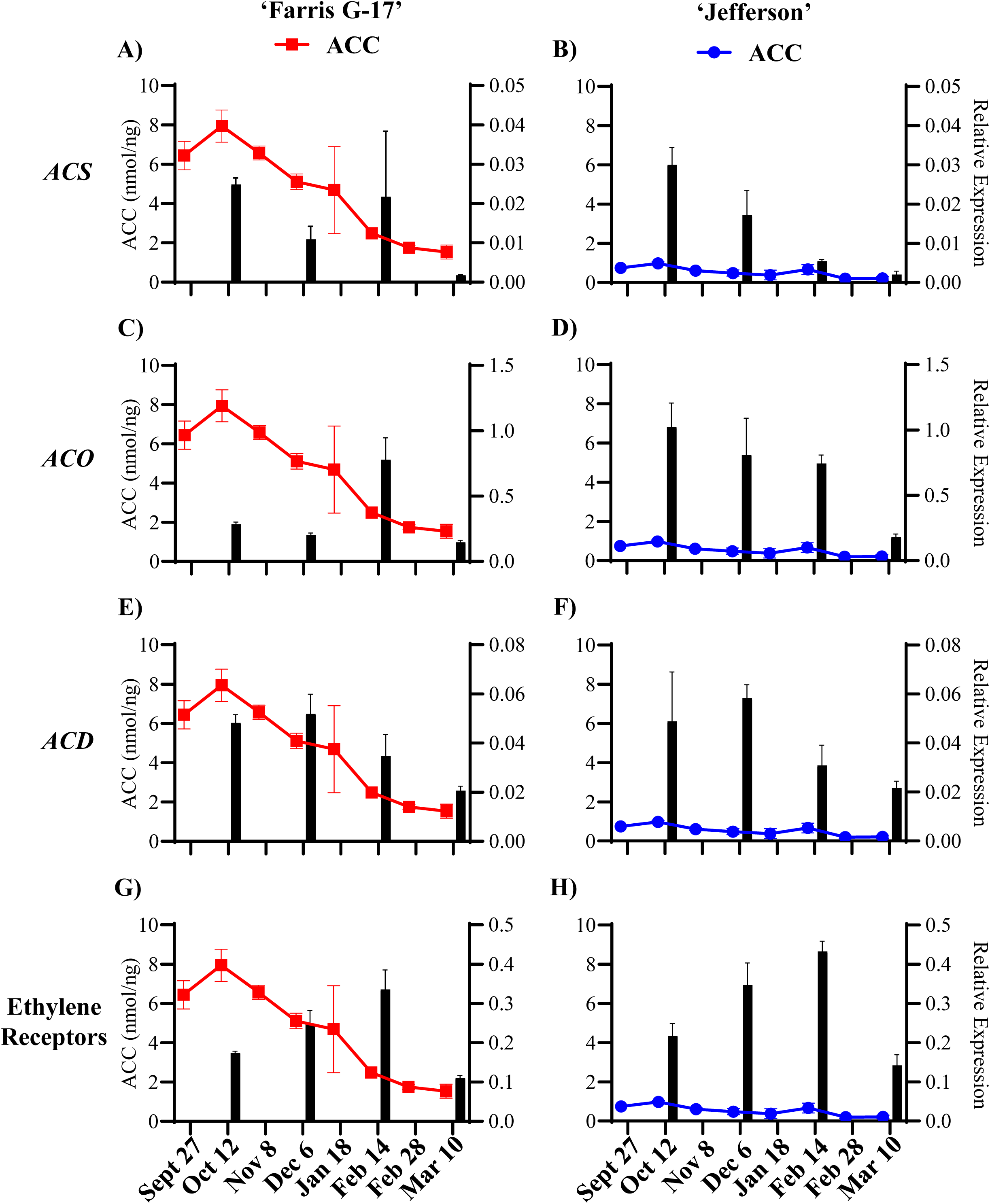
Total ACS, ACO, ACD, and ethylene relative gene expression as well as ACC levels of hazelnut catkins during the 2021/2022 season. A) The average relative gene expression of pooled ACS genes (ACS2, ACS3, ACS11) as well as ACC (nmol/g) in ‘Farris G-17′ catkins (n=2). Error bars represent standard deviation. B) The average relative gene expression of pooled ACS genes (ACS2, ACS3, ACS11) as well as ACC (nmol/g) in ‘Jefferson′ catkins (n=3). Error bars represent standard deviation. C) The average relative gene expression of pooled ACO genes (ACO2, ACO5, ACO6, ACS11) as well as ACC (nmol/g) in ‘Farris G-17′ catkins (n=2). Error bars represent standard deviation. D) The average relative gene expression of pooled ACO genes (ACO2, ACO5, ACO6, ACS11) as well as ACC (nmol/g) in ‘Jefferson′ catkins (n=3). Error bars represent standard deviation. E) The average relative gene expression of ACD as well as ACC (nmol/g) in ‘Farris G-17′ catkins (n=2). Error bars represent standard deviation. F) The average relative gene expression of ACD as well as ACC (nmol/g) in ‘Jefferson′ catkins (n=3). Error bars represent standard deviation. G) The average relative gene expression of pooled ethylene receptor genes (ETR1, ETR2, EIN4) as well as ACC (nmol/g) in ‘Farris G-17′ catkins (n=2). Error bars represent standard deviation. H) The average relative gene expression of pooled ethylene receptor genes (ETR1, ETR2, EIN4) as well as ACC (nmol/g) in ‘Farris G-17′ catkins (n=3). Error bars represent standard deviation.

Total *ACS* relative gene expression appears to be approximately equal between ‘Farris G-17′ and ‘Jefferson’ (Figure 13A, B). *ACS3* is the predominant *ACS* in catkins across all timepoints, in both accessions (Figure 14A, B). Total *ACO* relative gene expression levels seem to differ between accessions on October 12^th^ and December 6^th^, but appear to be the same thereafter (Figure 13C, D). These discrepancies are due to high relative expression of *ACO2* in ‘Jefferson’ (Figure 14C, D). The pattern and magnitude of total *ACD* relative gene expression is approximately equal between ‘Farris G-17′ and ‘Jefferson’ (Figure 13E, F). *ACD* relative gene expression appears to be high on October 12th and December 6th, and decreases on February 14th and March 10th, along with ACC levels (Figure 13E, F). The pattern and magnitude of total ethylene receptor relative gene expression is approximately equal between ‘Farris G-17′ and ‘Jefferson’ (Figure 13G, H). Ethylene receptors’ relative gene expression appears to increase from October 12^th^ to February 14^th^, then decrease on March 10^th^ (Figure 13G, H). These trends appear to be consistent for each individual ethylene receptor (Figure 14E, F).

**Figure 14.**
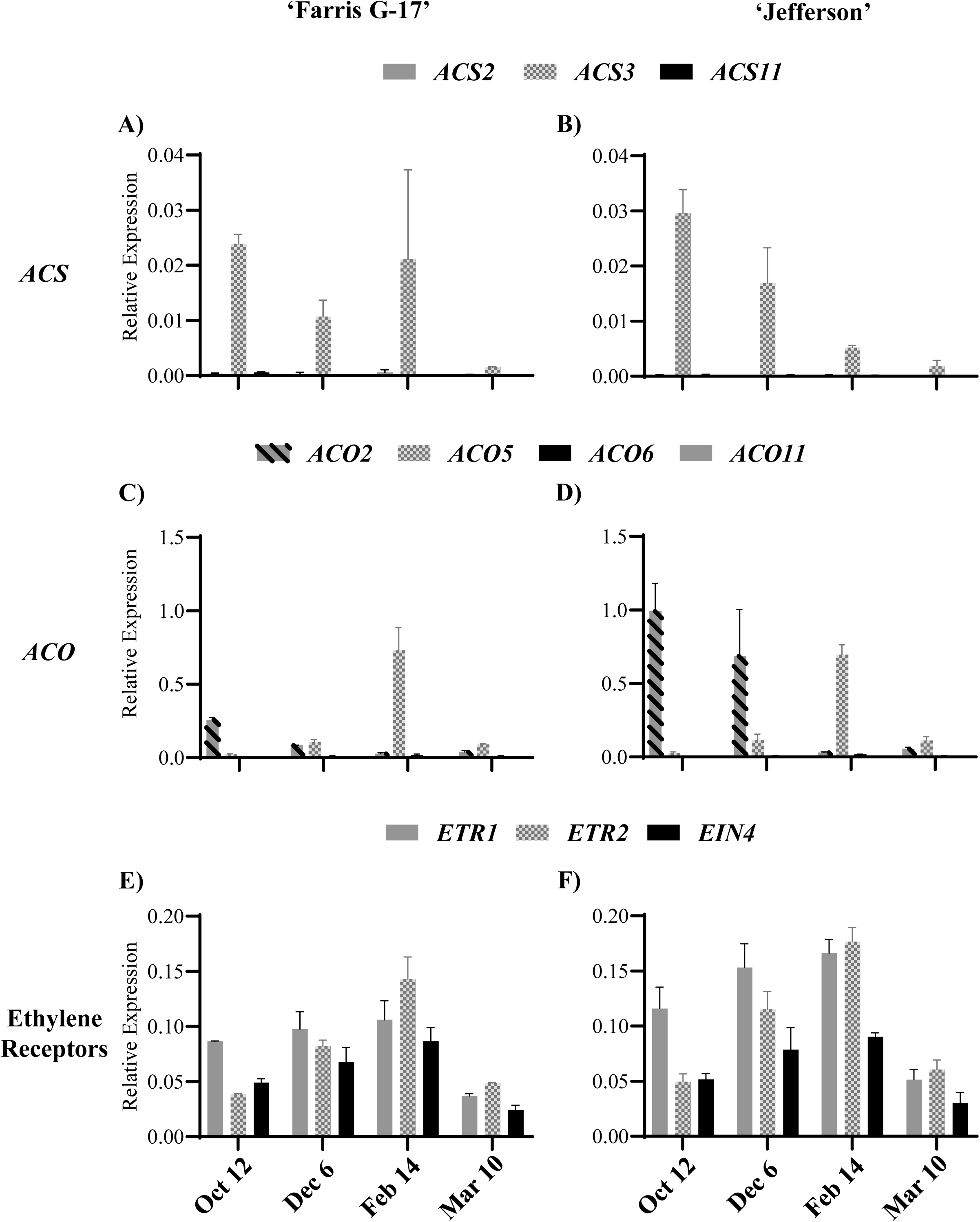
ACS, ACO and ethylene receptor relative transcript abundance in hazelnut catkins during the 2021/2022 season. A) The average relative gene expression of ACS2, ACS3 and ACS11 in ‘Farris G-17′ catkins (n=2). Error bars represent standard deviation. B) The average relative gene expression of ACS2, ACS3, ACS11 in ‘Jefferson′ catkins (n=3). Error bars represent standard deviation. C) The average relative gene expression of ACO2, ACO5, ACO6 and ACS11 in ‘Farris G-17′ catkins (n=2). Error bars represent standard deviation. D) The average relative gene expression of ACO2, ACO5, ACO6 and ACS11 in ‘Jefferson′ catkins (n=3). Error bars represent standard deviation. E) The average relative gene expression of ETR1, ETR2, EIN4 in ‘Farris G-17′ catkins (n=2). Error bars represent standard deviation. F) The average relative gene expression of ETR1, ETR2, EIN4 in ‘Farris G-17′ catkins (n=3). Error bars represent standard deviation.

Across the season, ACC levels appear to be correlated with *ACS* relative gene expression (Figure 13A,B). Indeed, *ACS* relative gene expression and ACC have a positive Pearson Correlation Coefficient of 0.626 in ‘Farris G-17′ and 0.795 in ‘Jefferson’. However, ACC patterns in ‘Farris G-17’ appear to be more closely related to the ratio of *ACS* to *ACO* relative gene expression (Figure 15A,B). The *ACS*/*ACO* relative gene expression ratio has a Pearson Correlation Coefficient of 0.999 in ‘Farris G-17’ and 0.549 in ‘Jefferson’ to ACC. Interestingly, an increased *ACS*/*ACO* relative gene expression ratio in ‘Farris G-17’ is associated with increased ACC levels compared to ‘Jefferson’ (Figure 15A,B). Taken together, this implies that different transcriptional regulation of *ACO* by ‘Farris G-17’ and ‘Jefferson’ may contribute to differences in ACC accumulation and ethylene biosynthesis.

**Figure 15.**
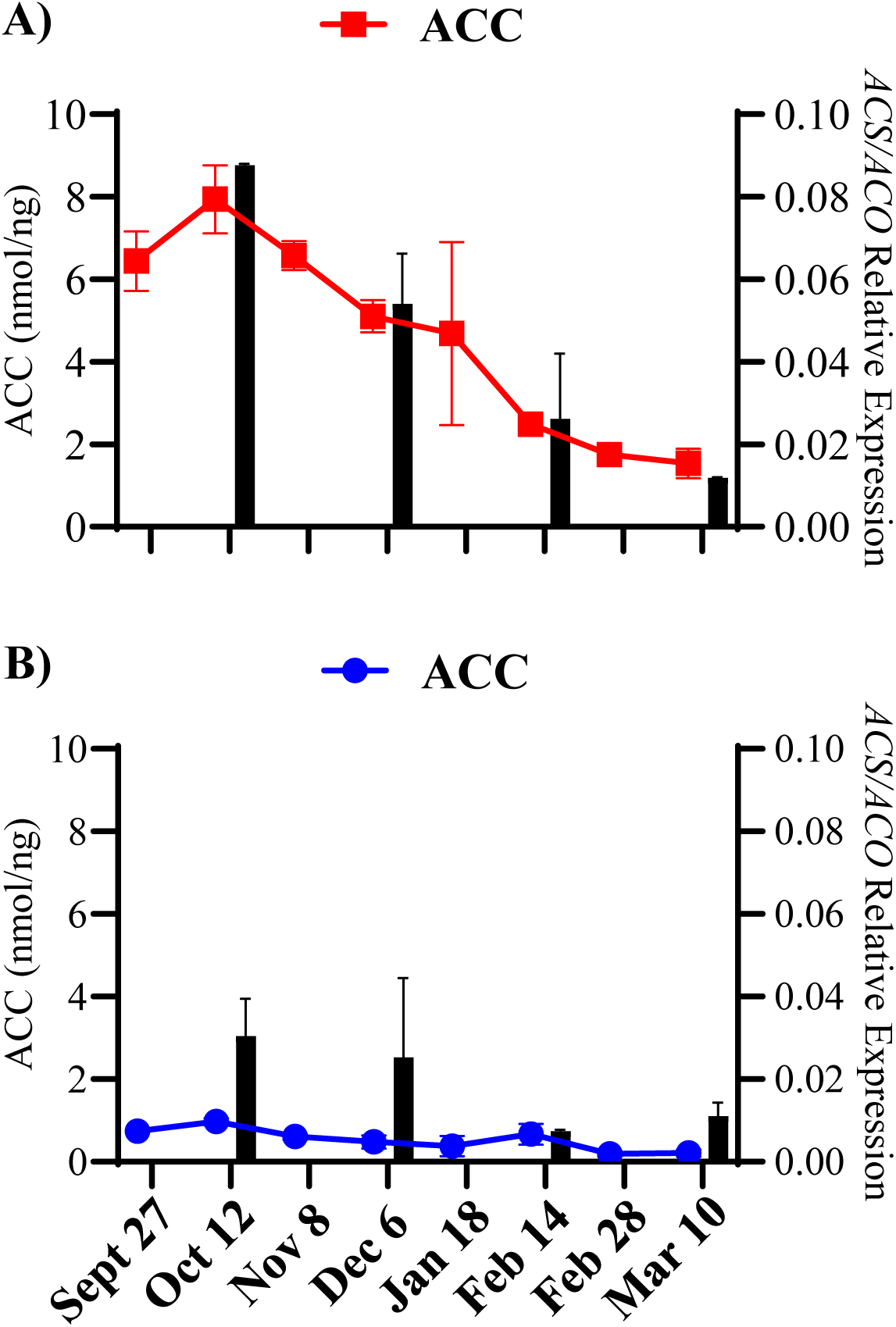
The ratio of total ACS to total ACO relative gene expression as well as ACC levels of hazelnut catkins during the 2021/2022 season. A) The average ratio of relative gene expression of pooled ACS genes (ACS2, ACS3, ACS11) to the relative gene expression of pooled ACO genes (ACO2, ACO5, ACO6, ACS11) as well as ACC (nmol/g) in ‘Farris G-17′ catkins (n=2). Error bars represent standard deviation. B) The average ratio of relative gene expression of pooled ACS genes (ACS2, ACS3, ACS11) to the relative gene expression of pooled ACO genes (ACO2, ACO5, ACO6, ACS11) as well as ACC (nmol/g) in ‘Farris G-17′ catkins (n=2). Error bars represent standard deviation.

### 2.5 Summary of Hormonal Trends

To view the trends of hormones in relation to each other, the average values of ABA, pooled GAs, IAA, pooled active CTKs, and ACC of all trees were adjusted to be at the same scale (Figure 16). Generally, ABA, IAA, and ACC are high during paradormancy while GA and CTK levels are low. In endodormancy, ABA peaks while IAA and ACC decrease (Figure 16). GA and CTK steadily increase in endodormancy (Figure 16). In ecodormancy, ABA decreases as well as ACC, and both reach their minimal levels (Figure 16). IAA remains steady from endodormancy to ecodormancy at its minimal levels, while GA and CTK reach and maintain their peak levels throughout ecodormancy (Figure 16).

**Figure 16.**
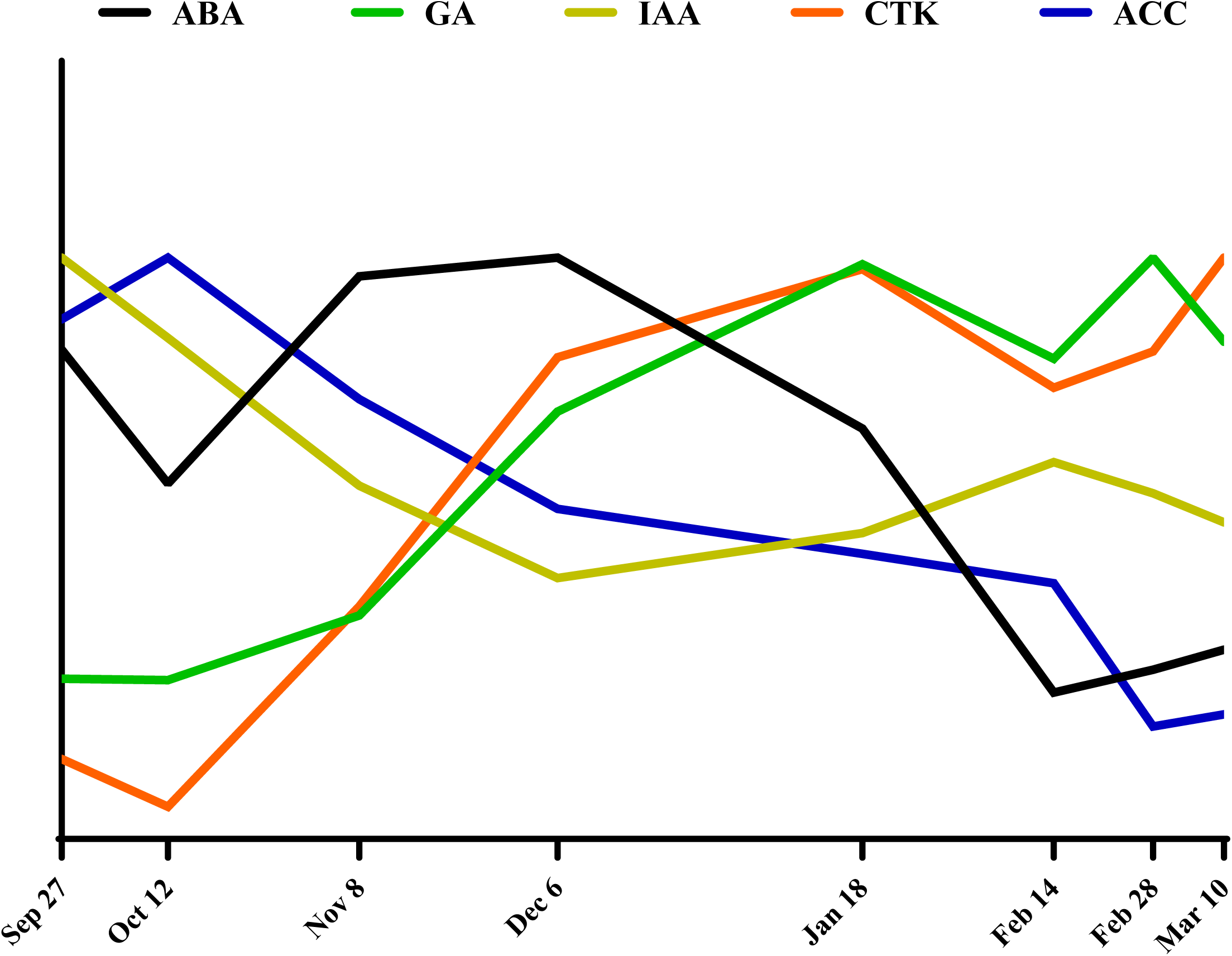
Representative hormone trends in dormant hazelnut catkins during the 2021/2022 season. Shown above are the representative trends of bioactive ABA, pooled GA, IAA, pooled active CTKs (CTK), and ACC in dormant hazelnut catkins during the 2021/2022 winter season. Representative trends for each hormone are the average values of all 7 trees included in the study. Hormone levels (nmol/g) were multiplied by a factor so that each hormone has the same maximum value of 1.5. “para” = paradormancy, “endo” = endodormancy, “eco” = ecodormancy.

### 2.6 Hormone Ratios

Due to well documented control of seed dormancy through the ABA/GA ratio, the ABA/GA ratio is presented here (Figure 17A). During paradormancy the ABA/GA ratio is erratic, however during endodormancy the ratios steadily decrease and appear to remain low during ecodormancy (Figure 17A). Looking at the inverse ratio, GA/ABA, can reveal trends in ecodormancy that would otherwise be masked. Indeed, the GA/ABA ratio increases during ecodormancy (Figure 17B). It appears there is some segregation between varieties in the GA/ABA ratio during ecodormancy, with ‘Farris 182’ having a higher GA/ABA ratio than ‘Farris G-17′ and ‘Jefferson’ (Figure 17B). ‘Farris G-17′ and ‘Jefferson’ have a similar GA/ABA ratio while *C. americana* has the lowest GA/ABA ratio (Figure 17B).

**Figure 17.**
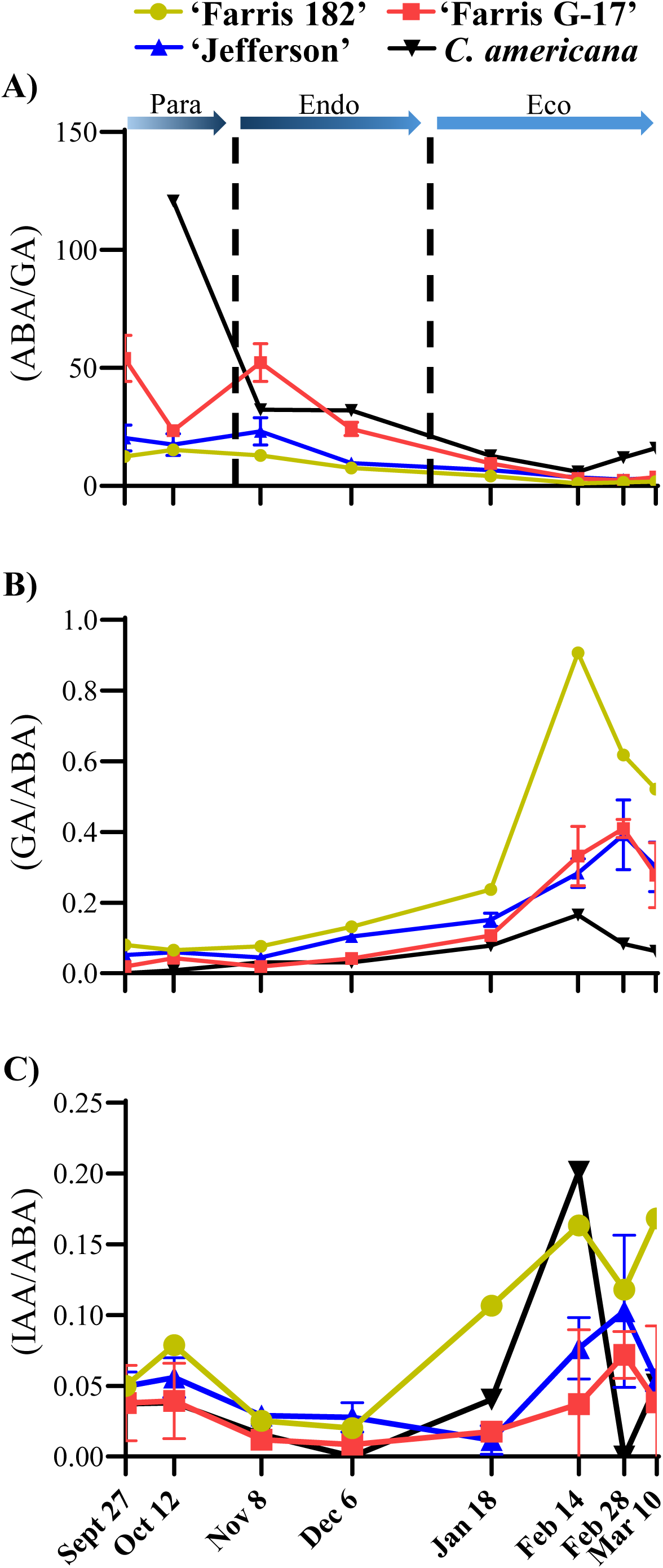
ABA/GA, GA/ABA, and IAA/ABA ratios in dormant hazelnut catkins during the 2021/2022 season. A) The ABA/GA ratio in ‘Farris 182’ (n=1), ‘Farris G-17’ (n=2), ‘Jefferson’ (n=3), and C. americana (n=1) catkins during the 2021/2022 season. Error bars indicate standard deviation. Vertical dashed lines indicate likely dormancy transitions. Colour gradients within arrow bars reflect the gradual transitions between dormancy states likely to occur. “para” = paradormancy, “endo” = endodormancy, “eco” = ecodormancy. B) The GA/ABA ratio in ‘Farris 182’ (n=1), ‘Farris G-17’ (n=2), ‘Jefferson’ (n=3), and C. americana (n=1) catkins during the 2021/2022 season. Error bars indicate standard deviation. Vertical dashed lines indicate likely dormancy transitions. Colour gradients within arrow bars reflect the gradual transitions between dormancy states likely to occur. “para” = paradormancy, “endo” = endodormancy, “eco” = ecodormancy. C) The IAA/ABA ratio in ‘Farris 182’ (n=1), ‘Farris G-17’ (n=2), ‘Jefferson’ (n=3), and C. americana (n=1) catkins during the 2021/2022 season. Error bars indicate standard deviation. Vertical dashed lines indicate likely dormancy transitions. Colour gradients within arrow bars reflect the gradual transitions between dormancy states likely to occur. “para” = paradormancy, “endo” = endodormancy, “eco” = ecodormancy.

Although the ratio of ABA to auxin has not been proposed to primarily regulate dormancy, their ratio has been measured in dormant female/vegetative buds of hazelnut before and so is also presented here (Figure 17C). The IAA/ABA ratio decreases from paradormancy into endodormancy, and spikes in ecodormancy, leading toward bloom (Figure 17C). The IAA/ABA ratio is relatively similar between all accessions during paradormancy and endodormancy, but highly variable during ecodormancy (Figure 17C).

## Chapter 3 Discussion

### 3.1 ABA

Generally, the bioactive ABA content of hazelnut catkins, measured in the present study, is similar to observations made in sweet cherry, pear, grape, apricot, and peach (Götz & Chmielewski, 2023; Chmielewski et al., 2018; Vimont et al, 2021; Gao et al., 2021; Ito et al., 2021; Li et al., 2018; Or et al, 2000; Zheng et al., 2015; Zhang et al., 2018; Liu & Islam et al., 2021). ABA levels increase during the transition from paradormancy to endodormancy, peak during endodormancy and decrease in ecodormancy (Figure 7E). This result provides further evidence for the primary role of ABA in promoting and maintaining bud endodormancy.

Between accessions, there were great differences in DPA content, but similar amounts of bioactive ABA (Figure 7A,B,C,D,F). An increased level of DPA indicates higher flux through the ABA biosynthetic pathway. This suggests that despite higher ABA biosynthesis inherent to different accessions, such as *C. americana*, bioactive ABA levels are tightly regulated. Bioactive ABA content appears to be primarily regulated through ABA catabolism, opposed to ABA conjugation, as accessions have very different DPA content across the season, but relatively similar ABA-GE content (Figure 7A,B,C,D). Despite tight regulation of ABA content through catabolism, it is interesting to note that ABA-GE accumulates at the expense of ABA in ecodormancy, whereas DPA does not (Figure 7A,B,C,D). The apparent negative correlation between ABA and ABA-GE, and the relative steady levels of DPA, indicate that ABA conjugation could be the main mechanism to regulate active ABA content during ecodormancy. Indeed, it is thought that ABA-GE formation is the primary form of ABA inactivation in Paris polyphylla seed dormancy release (Zheng et al., 2023). A similar trend is observed in grape buds, where ABA-GE was proposed to accumulate at the expense of ABA, while PA and DPA content remained constant (Zheng et al., 2015). Hints of this type of fine regulation were also found at the transcriptional level in Japanese apricot (Zhang et al., 2018). Zhang et al. (2018) found increased transcription of *UGT71B6*, a gene that glycosylates ABA, during the endo to ecodormant transition, indicating increased glycosylation may be key to inactivating ABA in ecodormancy. However, there is conflicting evidence for this kind of fine regulation of ABA in sweet cherry flowers. Götz & Chmielewski (2023) and Chmielewski et al. (2018) found ABA-GE accumulated at the expense of ABA in ecodormancy of cherry, but no such relationship was found by Vimont et al (2021). Taken together, it is clear that ABA catabolism and glycosylation play an important role in the regulation of ABA content in dormant flowers, although further investigation is required to determine the significance of ABA glycosylation in the regulation of ecodormancy.

### 3.2 GA

GA has been measured in sweet cherry, pear, grape, apricot, and peach (Vimont et al., 2021; Chengguo et al., 2004; Ito et al., 2021; Tamura et al., 2002; Zheng et al., 2018; Zhang et al., 2018; Wen et al., 2016; Lin et al., 2018; Liu & Islam et al., 2021). Usually, only bioactive gibberellins (GA_1_, GA_3_, GA_4_, or GA_7_) have been targeted in these species. GA is generally low in endodormancy and increases toward ecodormancy (Vimont et al., 2021; Chengguo et al., 2004; Wen et al., 2016; Lin et al., 2018; Liu & Islam et al., 2021). Although, a study in pear only showed increases during ecodormancy, and a study in apricot only showed an increase immediately before bloom (Ito et al., 2021; Zhang et al., 2018). In the present study, bioactive gibberellins (GA_1_, GA_2_, GA_4_, or GA_7_) could be sporadically detected, but were below the limit of quantification. However, precursors to GA_1_ (GA_19_, GA_44_, GA_53_) and the immediate catabolite of GA_1_ (GA_8_) could be measured. Previous hormone profiling in hazelnut plantlets has shown similar results, a failure to detect GA_1_, along with low quantities of GA_1_ precursors and catabolites (Erland et al., 2020). Bioactive GAs were also not detected in the development of sweet cherry buds by Chmielewski et al. (2018) or in peach by Liu & Islam et al. (2021). The inability to detect bioactive GAs during dormancy could be reflective of the minute amounts of GA needed to elicit a response, or that there is a fast flux through the GA pathway and products are rapidly catabolized after performing their role. Regardless, it is clear that future studies should include GA metabolites in addition to bioactive GAs to capture as much information about this important hormone as possible. In the present study, GA content was low during paradormancy, steadily increased during endodormancy, and remained high during ecodormancy (Figure 8E). The increase in GA content from endodormancy toward ecodormancy observed in the present study, matches well with observations made in sweet cherry and apricot (Vimont et al., 2021; Chengguo et al., 2004; Wen et al., 2016; Lin et al., 2018).

In the literature there is mixed evidence for the abundance of GA in paradormant flower buds. One study in sweet cherry, one study in pear, and one study in grape show high GA during paradormancy (Chengguo et al., 2004; Ito et al., 2021; Zheng et al., 2018) while another study in sweet cherry and a study in apricot show low GA during paradormancy (Vimont et al., 2021; Wen et al., 2016). The results of the present study agree with the latter, that GA is low during paradormancy (Figure 8E). In the literature there is also mixed evidence for the abundance of GA throughout ecodormancy. One study in sweet cherry and a study in pear show continual increases in GA during ecodormancy (Chengguo et al., 2004; Ito et al., 2021) while another study in sweet cherry shows a decrease in GA during ecodormancy (Vimont et al., 2021). The results of the present study agree with neither trend, showing steady GA content throughout ecodormancy (Figure 8E).

#### Auxin

Endogenous auxin content has been measured in dormant flower buds of pear and apricot, as well as hazelnut female/vegetative buds (Ito et al., 2021; Zhang et al., 2018; Rodríguez & Sánchez□Tamés, 1986). Bioactive auxin, IAA, was the only auxin metabolite measured in these studies. In pear, IAA content was low throughout dormancy, spiking at or just before bloom (Ito et al., 2021). In apricot, IAA content was relatively steady in endo and ecodormancy, spiking at bud burst (Zhang et al., 2018). In Spain, IAA content in hazelnut female/vegetative buds decreased from October to November, then steadily increased toward bloom (Rodríguez & Sánchez□Tamés). The results of the present study generally agree with observations in pear and apricot, with IAA content remaining relatively constant throughout endodormancy and ecodormancy (Figure 9E). However, a spike just before bloom was not noticed in the present study, as it was in pear (Ito et al., 2021). Similar to hazelnut buds in Spain, IAA content decreased from October into November, although the present study did not observe an increase in IAA content afterwards (Rodríguez & Sánchez□Tamés).

High levels of IAA at the onset of paradormancy may be compared to IAA in the regulation apical dominance of axillary buds. Paradormancy is like axillary dormancy in the sense that paradormant buds are not regulated primarily by endogenous factors, but by growth factors produced from external organs (Lang et al., 1987). Thus, increased IAA may play a role in the establishment and/or maintenance of paradormancy, although further investigation is required to confirm this hypothesis. Alternatively, since IAA is also known to promote growth, increased IAA in paradormancy could be reflective of the increased ability of paradormant buds to bloom under forcing conditions.

Interestingly, there are differences in IAA-Asp patterns between accessions. In ‘Jefferson’ and *C. americana*, IAA-Asp seems to correlate with IAA throughout the season (Figure 9C,D). In ‘Farris 182’ and ‘Farris G-17′, however, IAA-Asp is generally not detected in endodormancy and ecodormancy (Figure 9A,B). Since conjugation to Asp is currently considered to be the first step in IAA catabolism, decreased IAA-Asp in ‘Farris 182’ and ‘Farris G-17’ may indicate that less IAA is being catabolized to maintain free IAA levels. Alternatively, increased IAA-Asp in ‘Jefferson’ and *C. americana* may simply indicate higher flux through the biosynthetic pathway. It is impossible to know which, if either, is the case without further investigation. One intriguing idea is that the difference may be due to different Trp pools. Trp is a precursor to not only IAA, but also indoleamines (Murch et al., 2000). Indoleamines have been implicated in cold tolerance, and it is possible that ‘Farris G-17′ is shuffling more Trp toward indolamine biosynthesis in an attempt to combat cold stress during the winter period (Ayyanath et al., 2021). As a result, less free IAA may be produced, and IAA levels may be preserved via reduced catabolism. A more comprehensive profile of auxin metabolites and indolamines would be necessary to test this hypothesis, which remains highly speculative.

### 3.3 CTK

The endogenous levels of CTK in deciduous tree bud dormancy have not been extensively investigated thus far. Only recently, active CTK levels have been measured in dormant Japanese pear buds including tZ and iP type CTKs (Ito et al., 2021). In pear, the tZ type CTKs were low throughout endodormancy and ecodormancy, and sharply increased immediately before bloom. In contrast, iP type CTKs increased during the transition from endodormancy to ecodormancy, peaking around the transition and decreased thereafter. Thus, in Japanese pear, it appears as though iP was correlated with endodormancy transition and tZ was primarily found during ecodormancy release.

In the present study, CTK precursors (iPR, tZR, dhZR, and cZR), active CTKs (iP, tZ, dhZ, cZ) and O-glucoside conjugates (tZOG, cZOG) were measured, expanding the breadth of cytokinin metabolites measured during flower dormancy. Active CTK content was low during paradormancy, increased during endodormancy, and remained high in ecodormancy leading toward bloom (Figure 11E). This pattern is very different from observations in pear, which did not show consistently increasing CTK in endodormancy, or sustained CTK content in ecodormancy (Ito et al., 2021). tZ did not have a distinctive pattern to compare to that observed by Ito et al. 2021, however iP has a strikingly similar increase during the transition from endodormancy to ecodormancy, albeit without a following decrease in ecodormancy.

cZR and cZOG appeared to be relatively constant throughout the study compared to the other CTK metabolites (Figure 10 A-H). cZ has, for a long time been regarded as a mere catabolite of tRNA, with irrelevant CTK activity, due to its much lower activities compared to tZ in traditional growth assays (Schäfer et al., 2015). Constant levels of cZR type CTKs, as observed in the present study, would be consistent with this theory. Although, because cZ has lower activity than tZ and iP, it may play a specialized role in maintaining baseline CTK activity in tissues where growth has slowed or halted, such as dormant tissues (Gajdošová et al., 2011; Schäfer et al., 2015). Given the very low levels of active CTKs observed during paradormancy, and relatively high cZR content, it is possible that cZ is providing baseline CTK activity during this stage. Although, since cZ itself was not detected in the present study, this role for cZ is speculative.

### 3.4 Evaluation of ethylene biosynthesis in catkins

Endogenous ethylene biosynthesis has not yet been measured in dormant flowers of deciduous woody perennials. Recently, however, the endogenous levels of ACC, the immediate precursor to ethylene, have been measured in endodormant pear flowers (Gao et al., 2021). In Gao et al. 2021, ACC was measured in endodormant flowers from shoots exposed to artificial chilling, until the transition to ecodormancy. It was discovered that ACC was initially high in early endodormancy, decreased into mid endodormancy, and increased at endodormancy release. Because it is not feasible to measure ethylene levels in a field experiment, ACC was also used as a proxy for ethylene biosynthesis in the present study. The results of the present study partly agree with observations in pear, with ACC content decreasing during endodormancy. Contrary to pear, however, ACC continued to decrease during ecodormancy (Figure 29A). This ACC trend also conflicts with observations made in dormant seeds as ethylene generally increases with dormancy progression in seeds, working antagonistically to ABA (Corbineau et al., 2014). Indeed, ethylene applications to both dormant seeds and endodormant buds have been found to promote dormancy release, and ethylene signaling inhibitors have delayed dormancy release (Corbineau et al., 2014; Ophir et al., 2009).

If it is assumed that ACS regulates the rate-limiting step in ethylene biosynthesis, ACC can be used as a proxy for ethylene biosynthesis. Assuming this is true in the present study, it could be proposed that ethylene biosynthesis is greatest at paradormancy and decreases toward ecodormancy release. A positive correlation between ACC and *ACS* relative gene expression across the season, as observed, would support this interpretation. This interpretation would also be supported by decreasing *ACO* relative gene expression observed in ‘Jefferson’, as decreasing *ACO* expression would also suggest decreasing ethylene biosynthesis.

Interestingly, while differences in all other hormone levels between early and late-blooming accessions were subtle, early-blooming ‘Farris G-17′ had significantly more ACC than late-blooming ‘Jefferson’ during most of the study. It could be therefore be inferred that ‘Farris G-17’ produces more ethylene than ‘Jefferson’ throughout the season. However, it is difficult to support this assumption since differences in ACC levels between accessions cannot be explained by differing *ACS* or *ACD* relative gene expression. Neither could differences in ethylene receptor relative gene expression compensate for differences in ACC levels, as they are also very similar between accessions. Instead, the discrepancy appears to be explained by *ACS*/*ACO* relative gene expression ratio. The *ACS/ACO* relative gene expression ratio is much higher in ‘Farris G-17’ than ‘Jefferson’ and is highly correlated with ACC levels in ‘Farris G-17’. This finding suggests ACC may be accumulating in ‘Farris G-17’ due to decreased ACO availability compared to ‘Jefferson’ and therefore that ACO may be regulating the rate-limiting step in ethylene biosynthesis in dormant hazelnut catkins of ‘Farris G-17’. This would be contrary to the general assumption that ACS regulates the rate-limiting step of ethylene biosynthesis, but not unprecedented. ACO has been found to regulate the rate-limiting step in ethylene biosynthesis in post-climacteric tomato fruit ripening (Van de Poel et al., 2012). There is also evidence for this kind of regulation of ethylene biosynthesis by ACO in the petioles of flooded tomato, the formation of tension wood in poplar, cotton fibre cell elongation, and sex determination in cucumber flowers (English et al., 1995; Andersson-Gunnerås et al., 2003; Shi et al., 2006; Chen et al., 2016). If ACO is being regulated in such a way in ‘Farris G-17’ catkins, it would suggest ACC is accumulating in ‘Farris G-17’ due to decreased ethylene biosynthesis compared to ‘Jefferson’. However, it is impossible to determine this without directly measuring ethylene production of the catkins. Additionally, there are other factors that could explain the discrepancy in ACC accumulation between cultivars that were not investigated in the present study, such as ACC conjugation or post-translational modifications to ethylene metabolic enzymes. Therefore, further investigation into the regulation of ethylene biosynthesis during catkin dormancy is needed to determine which accession truly produced more ethylene, if either. Even though there is doubt as to how to interpret increased ACC in ‘Farris G-17’, it remains likely that ethylene biosynthesis decreases from paradormancy through ecodormancy in the present study, since both ACS and ACO gene expression levels tend to decrease along with ACC as dormancy progresses.

#### Hormone Ratios

The ratio of antagonistic hormones, rather than absolute levels of hormones, can sometimes be more informative. For example, it is widely recognized that the ABA/GA ratio regulates seed dormancy (Ali et al., 2022). The ratio of antagonistic hormones is typically not reported in dormant flower buds, save for a couple studies which presented the ABA/GA ratio and one study which measured the IAA/ABA ratio in dormant hazelnut female flowers/vegetative buds (Rodríguez & Sánchez□Tamés, 1986). The ABA/GA ratio measured in the present study is similar to the ratios reported in sweet cherry and apricot, supporting the notion that GA behaves antagonistically to ABA as a growth promoter during dormancy regulation (Figure 17A) (Chengguo et al., 2004; Zhang et al., 2018). Compared to the ABA/GA ratio, the IAA/ABA ratio is more complex. IAA/ABA ratios are high during paradormancy, decrease slightly in endodormancy, and appear to increase in ecodormancy (Figure 17C). The increasing IAA/ABA ratio from endodormancy to ecodormancy match the trends observed in hazelnut female flower/vegetative buds by Rodríguez & Sánchez□Tamés (1986). The increasing IAA/ABA ratio from endodormancy to ecodormancy implies IAA and ABA act antagonistically during ecodormancy. Thus, IAA may play a role in promoting ecodormancy release. This would be consistent with observations made by Ito et al. (2021), who observed spikes of auxin just before bloom in pear.

#### Recommendations for the application of hormones to extend dormancy in hazelnut catkins

The application of ABA analogs to dormant grape buds in the fall has successfully delayed bloom in the subsequent spring in grape. Since patterns for endogenous ABA content are similar between dormant hazelnut catkins and grape buds, it is reasonable to suggest ABA analogs may have the same effect if applied to hazelnut trees. Interestingly, ABA applications appear to be more effective in extending dormancy if applied during early endodormancy and are not effective when applied during late endodormancy or ecodormancy in grape, peach, pear, and sweet cherry flower buds (Bowen et al., 2016; Parker et al., 2012; Tamura et al., 2002; Vimont et al., 2021). Furthermore, ABA has been shown to delay endodormancy release in a dose-dependent manner in grape (Zheng et al., 2015). Decreasing effectiveness of ABA applications during endodormancy progression, and increased effectiveness with higher doses of ABA, may be due to the relative levels of antagonistic hormones, such as GA or CTK. Thus, the ratio of ABA to antagonistic hormones could be key for regulating flower bud dormancy, as it is in seed dormancy. Since ABA analog applications were most effective delaying bloom when applied in the fall in grape, and since the ABA/GA ratio is highest in paradormancy and early endodormancy in the present study, it is recommended that ABA analogs also be applied to hazelnut trees in the fall.

## Conclusions

To conclude, by characterizing a vast hormone profile across an entire dormant season, the present study has provided new insights into the hormonal regulation of flower dormancy in deciduous woody perennials. Unexpectedly, CTK was found to have a pattern quite similar to that of GA across dormancy. Also, for the first time, ACC was observed to steadily decrease from paradormancy to ecodormancy release and large differences in ACC accumulation were noticed between early and late blooming accessions. This pattern could reflect a significant role for ethylene in the regulation of hazelnut catkin dormancy, but more studies are required to understand the nature of ethylene’s regulation. ABA and the ABA/GA ratio appeared to correlate will with dormancy depth, as seen in other crops. Thus, it is expected that applications of ABA analogs in the fall would delay bloom of hazelnut catkins in the following spring, as seen in grape (Bowen et al., 2016). Therefore, this work has not only provided a valuable resource for those studying flower dormancy, but has provided support for the application of ABA analogs to extend hazelnut catkin dormancy in Ontario. In the face of changing growing seasons due to climate change, it is our hope that this work will support sustainable agriculture around the globe as it has supported the hazelnut industry in Ontario.

